# The distribution of common-variant effect sizes

**DOI:** 10.1101/2020.09.19.304097

**Authors:** Luke Jen O’Connor

## Abstract

The genetic effect-size distribution describes the number of variants that affect disease risk and the range of their effect sizes. Accurate estimates of this distribution would provide insights into genetic architecture and set sample-size targets for future genome-wide association studies. We developed Fourier Mixture Regression (FMR) to estimate common-variant effect-size distributions from GWAS summary statistics. We validated FMR in simulations and in analyses of UK Biobank data, using interim-release summary statistics (max N=145k) to predict the results of the full release (N=460k). Analyzing summary statistics for 10 diseases (avg N_eff_=169k) and 22 other traits, we estimated the sample size required for genome-wide significant SNPs to explain 50% of SNP-heritability. For most diseases the requisite number of cases is 100k-1M, an attainable number; ten times more would be required to explain 90% of heritability. In well-powered GWAS, genome-wide significance is a conservative threshold, and loci at less stringent thresholds have true positive rates that remain close to 1 if confounding is controlled. Analyzing the shape of the effect-size distribution, we estimate that heritability accumulates across many thousands of SNPs with a wide range of effect sizes: the largest effects (at the 90^th^ percentile of heritability) are 100 times larger than the smallest (10^th^ percentile), and while the midpoint of this range varies across traits, its size is similar. These results suggest attainable sample size targets for future GWAS, and they underscore the complexity of genetic architecture.

## Introduction

Genome-wide association studies (GWAS) have detected thousands of associations with common diseases^1–4^. Despite their large size, these studies remain far from saturation: known loci explain well under 50% of SNP-heritability^5^, polygenic risk scores capture only a fraction of heritable risk^6^, and statistical power remains an obstacle for downstream analyses like fine mapping^7–9^ and heritability partitioning^10–12^.

If every human being were genotyped, would every genetic association be discovered? When is the point of diminishing returns? What fraction of the genome would be significantly associated with a disease, and how would the effect sizes of newly-discovered SNPs compare with those already discovered? These are questions about the genetic effect-size distribution, which describes how heritability accumulates across SNPs, from the few largest effects to the many smallest^1,13–15^. Various statistical methods estimate this distribution. Park et al.^16^ used cross validation to estimate the number of additional loci whose effect sizes are similar to those already discovered. Palla and Dudbridge^17^ fit a two-component mixture model, which assumes that all non-null loci have similar effect sizes (following a normal distribution). Several other methods fit similar models, with various objectives^13,18–21^. Moser et al.^22^ and Zhang et al.^14^ fit more sophisticated models, permitting a mixture of small- and large-effect SNPs. We previously quantified polygenicity without fitting a specific model^15^. A related but distinct objective is heritability partitioning, either across annotations^10–12^, regions^23,24^ or fine-mapped SNPs^25^. These methods have produced insights into genetic architecture, but critical questions remain unanswered about the future of genome-wide association studies and the distribution of heritability across the genome.

We introduce a method, Fourier Mixture Regression (FMR), to estimate genetic effect-size distributions. FMR operates on summary statistics, and it has the flexibility to fit almost any desired mixture model. Similar to LD score regression^10,26^, it reduces the estimation problem to linear regression, except that the regression coefficients quantify the heritability explained by the mixture components (e.g. large-effect vs. small-effect SNPs) rather than SNP annotations (e.g. coding vs. noncoding SNPs). To make this simplification, FMR relies on the Fourier transform, which maps between a time-varying signal (e.g. sound waves) and a distribution of frequencies (notes). We apply FMR to estimate the genetic effect size distribution of 32 diseases and complex traits.

## Definition of the effect-size distribution

Several studies^14,18,22^ have sought to estimate the distribution of causal, or joint-fit, effect sizes: the phenotypic effect of changing a single allele, without changing other alleles on the same haplotype. Causal effect sizes of individual SNPs are estimated in fine-mapping studies^7–9,25,27,28^. They differ from marginal effect sizes, or associations, which are duplicated across tag SNPs in high LD with one that is causal. Because of this duplication, the distribution of marginal effect sizes across SNPs does not determine the number of associated loci or the proportion of heritability that will be explained by GWAS, and it seems necessary to estimate the causal effect-size distribution, a challenging problem. However, for an individual SNP or locus, the power to detect it actually depends on its marginal effect size; therefore, for power analyses it would be preferable to address the challenge of marginal effect-size duplication without turning to causal effects.

We define the *heritability distribution of marginal effect sizes* (HDM), which describes the proportion of heritability explained by loci with various marginal effect sizes. For example, the median of the HDM is the effect size such that 50% of heritability is causally explained by SNPs with marginal effect sizes greater than the threshold. This definition sidesteps the problem of marginal effect-size duplication because the number of tag SNPs at a locus does not affect the heritability it explains. Therefore, estimates of the HDM can be used to predict the number of loci that will be detected in a GWAS and the heritability they will explain. The HDM is formally defined under a random-effects model for the purpose of estimation, but we actually report fractions of fixed-effect SNP-heritability^24^, which are more readily interpreted (see Methods).

An advantage of this definition is that the HDM is fundamentally easier to estimate than the distribution of causal effect sizes. If several associated SNPs are in perfect LD, it is impossible to tell based on GWAS data how many of them are causal; the distribution of causal effect sizes is unidentifiable without simplifying assumptions or heuristics. In contrast, the HDM is identifiable because the heritability explained by the LD block is determined by its marginal effect size (which is directly observed), and it makes no difference how many causal SNPs reside in that block. This distinction is relevant to real traits, as a single locus often harbors multiple causal SNPs^27^.

We measure effect sizes in per-s.d. units, i.e. in standard deviations of the phenotype per standard deviation of the genotype, rather than in per-allele units^5^. In per-s.d. units, common SNPs have larger effect sizes on average than rare SNPs^18^, and the squared marginal effect size of a SNP is equal to the heritability that it tags. It would also be of interest to quantify the HDM in per-allele units, but we use normalized units because they determine power in GWAS^29^ and because they can be interpreted as a fraction of heritability.

## Overview of Fourier Mixture Regression

In order to estimate the HDM, we developed Fourier mixture regression (FMR). FMR is a method-of-moments estimator for the mixing weights of an arbitrary finite mixture model for the HDM. Similar to stratified LD score regression^10,26^, it involves regressing a function of the GWAS summary statistics on SNP-specific scores computed from an LD reference panel. Whereas S-LDSC partitions heritability across annotations via multiple regression on annotation-specific LD scores^10^, FMR partitions heritability across mixture components via multiple regression on component-specific “Fourier LD scores” that depend on the effect-size distribution of each mixture component.

FMR relies on the Fourier transform, which maps between a time-varying signal (e.g. sound waves) and a distribution of frequencies (notes). The Fourier transform of a probability density function is called a characteristic function, denoted *ϕ*(*t*), and it fully characterizes a probability distribution. Here, characteristic functions enable us to convert a distribution over SNPs into a distribution over components of heritability, simply by taking a derivative. This convenient property comes from the characteristic function of the normal distribution, which is a bell curve (similar to the normal density function, except that *σ*^2^ appears in the numerator of the exponent). Let *X* ~ *N*(0, *σ*^2^). Its characteristic function is:

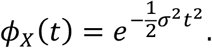

Taking the derivative, the variance parameter is pulled out:

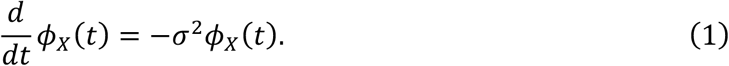

We can exploit this property to convert a distribution of effect sizes over SNPs into a distribution over components of heritability. Suppose that the effect size of a randomly-chosen SNP, *β*, follows a mixture of normal distributions with variance parameters 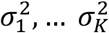 and mixture weights *a*_1_, … *a_K_*. The distribution of *β* over components of heritability is also a mixture of normal distributions, with the same variance parameters but with weights proportional to 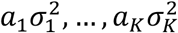. These two distributions have closely-related characteristic functions, and in particular, the latter is proportional to the derivative of the former:

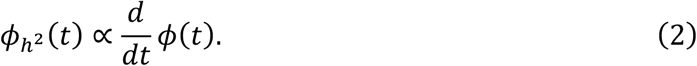

This observation leads to an estimation equation for the mixture weights of the HDM. Suppose the HDM is a mixture distribution whose mixture components have characteristic functions *ϕ*_1_, …, *ϕ_K_* and mixing weights *w*_1_, …, *w_K_*. Then the following equation allows us to estimate *w*:

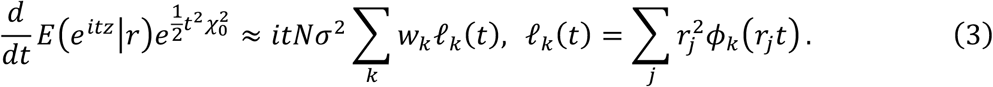

On the left-hand side, we have the derivative of the characteristic function of the GWAS Z score, *z*, with a noise correction. It depends on the sampling time *t* and the LD score regression intercept 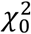, which accounts for population stratification and cryptic relatedness^26^. On the right-hand side, we have a linear combination of LD score-like terms ℓ*_k_*(*t*), one for each mixing component, with coefficients proportional to the mixing weights. These scores are a sum over SNPs in LD with the regression SNP, and they depend on the characteristic function of the mixing components. The sampling times are fixed. *N* is the effective GWAS sample size, *σ*^2^ is the average per-SNP heritability, and *i* is the imaginary unit (both sides of the equation are imaginary). We derive equation (3) in the Methods section. It relies on an LD approximation, roughly that LD between causal SNPs is either strong or weak.

We use 13 mixture components with variance parameters spanning a wide range of effect sizes (a factor of 4000 difference between the largest and smallest effect-size variances). The grid is centered on a trait-specific value that depends on its heritability and polygenicity (see Methods). This model can be thought of as an approximation to an infinite mixture model. Moser et al. used a mixture grid with 3 components and a factor of 100 difference between the smallest and largest-effect components^22^. A fixed-grid mixture model for a distribution of test statistics was also used by Stephens^30^.

In brief, FMR involves the following steps (see Methods for details): we specify a *K*-component mixture model and compute LD scores for the mixture components at *K* convenient sampling times. The LD scores are computed from a 1000 genomes reference panel^31^ with a correction for finite reference panel size. The HLA region is excluded. The variance of the mixture components must match the scale of the summary statistics; to avoid re-computing Fourier LD scores for every trait, the summary statistics are scaled to approximately match the Fourier LD scores. Then, we perform a weighted regression of the left-hand side of equation (1), evaluated at the *K* sampling times, on the corresponding LD scores multiplied by *it*. We also include moment equations from LDSC and LD 4th moments regression^15,26^. Regression weights are chosen using a heuristic. The regression is constrained to produce non-negative coefficients (this does not increase the computational complexity). We divide the coefficients by their sum to produce an estimate of *w*.

## Performance of FMR in simulations

We evaluated FMR in simulations using real LD from UK Biobank typed SNPs (M=455k SNPs) and a wide range of effect-size distributions. First, we performed simulations under the commonly-used point-normal model^17,20,32^, where a fraction of SNPs *M_c_/M* have causal effect sizes with normally distributed effects. Increasing *M_c_/M* shifts the HDM cumulative distribution function to the right: the shape of the distribution remains approximately constant, but most heritability is explained by fewer SNPs with larger effect sizes (Figure 1a-c). We applied FMR to summary statistics generated under these models at *N* = 460*k* and *h*^2^ = 0.1 (similar to UK Biobank; see below). Our method produced unbiased and robust estimates of the HDM across all values of *π_PN_* (Figure 1a-c).

**Figure 1:**
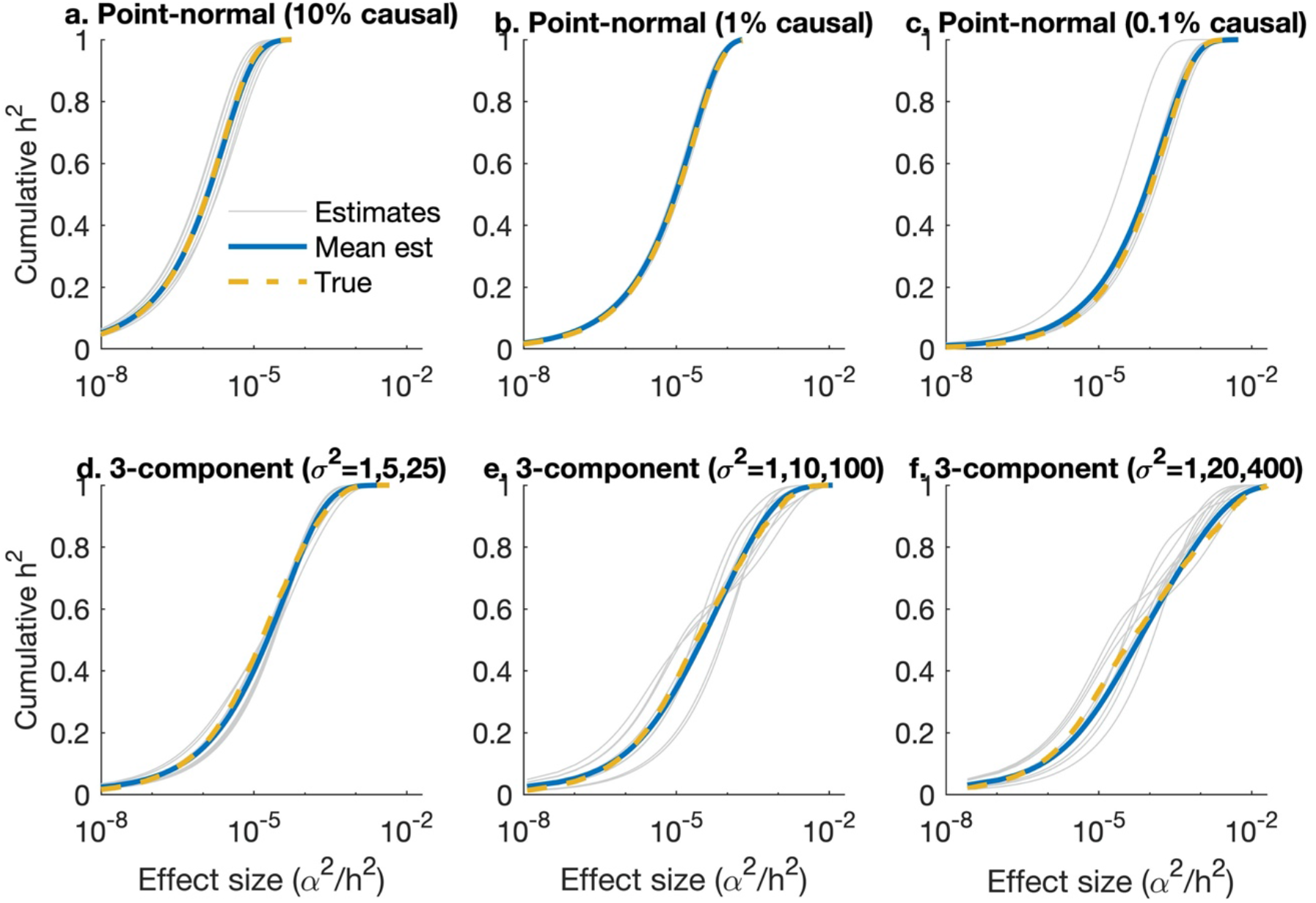
Performance of Fourier Regression in well-powered simulations (N=460k) with real LD. We show the true HDM, the mean estimated HDM (across 100 replicates), and ten individual HDM estimates (chosen at random). (a-c) Point-normal genetic architecture with different proportions of causal SNPs. (d-f) Gaussian mixture models with small-, medium, and large-effect SNPs. Each component explains 1/3 of heritability, and the relative effect size variance of the large vs. small-effect component is 25x, 100x and 400x in panels d-f respectively.

We simulated more complicated genetic architectures involving a mixture of small-, medium- and large-effect SNPs. Each non-null mixture component explained one third of heritability, and the total fraction of causal SNPs was approximately 1%. We varied the relative effect size of the three components such that the variance of the large-effect component was 25x, 100x, or 400x the variance of the small-effect component (5x, 10x and 20x the variance of the medium-effect component; Figure 1d-f). This change affects the slope of the HDM cumulative distribution function: under the 400x model (Figure 1f), the CDF has a shallow slope, and heritability is dispersed across a wide range of squared effect sizes (spanning ^~^5 orders of magnitude). In contrast, in Figure 1a-c, the slope is steep, and most heritability is explained by SNPs with similar effect sizes. We applied FMR to summary statistics generated under these genetic architectures and determined that it produces approximately unbiased estimates, though noisier than under the point-Normal model (Figure 1d-f). We note that the FMR model is not tailored to these specific simulations; in particular, the variance parameters of the FMR mixture components are not equal to those of the generative model.

We performed simulations at smaller GWAS sample sizes under the three point-Normal models. At small sample size (*N* = 50k, *h*^2^ = 0.1), FMR produced biased and noisy estimates (Figure S1a-c). At moderate sample size (*N* = 145k, similar to the UK Biobank interim release), FMR produced reliable estimates at *M_c_/M* ≤ 0.01 but biased and noisy estimates at *M_c_/M* = 0.1 (Figure S1d-f). It produced reliable estimates at N=460k (Figure 1). We identified an indicator for whether sample size was adequate: the Z score of the mean of the HDM (*Z*_*LD*4*M*_), which is estimated as a preprocessing step using LD 4^th^ moments regression^15^ (see Methods). We found that FMR is almost always well-powered when *z*_*LD*4*M*_ > 2, and in analyses of real phenotypes below, we exclude traits that do not meet this threshold.

## Performance of FMR predictions in UK Biobank

We applied FMR to summary statistics for 24 complex traits from the UK Biobank interim release (maximum N=145k) in order to predict the results of the full UK Biobank release (maximum N=460k). We predicted the number of loci that would be genome-wide significant (*M_GWAS_*) and the proportion of heritability they would explain 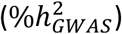, and we compared these predictions with observed values, which were much larger at N=460k than at N=145k (Figure S2). FMR predictions were approximately unbiased (mean 0.42 vs. 0.43 for 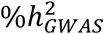, 786 vs. 731 for *M_GWAS_*), and they were highly correlated with observed values (Figure 2; r^2^=0.94 for 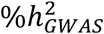, r^2^=0.92 for *M_GWAS_*), indicating that FMR can be used to project the results of future GWAS.

**Figure 2:**
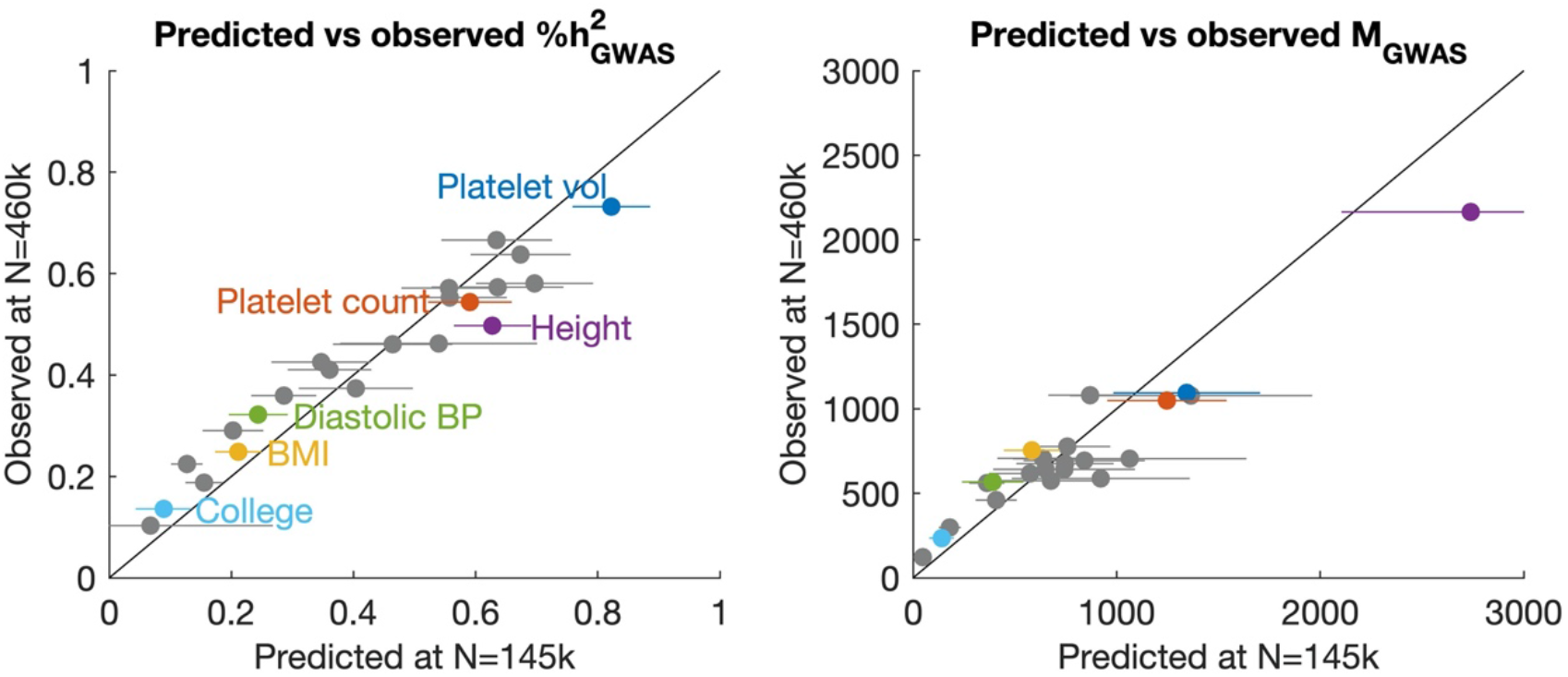
Predicted vs. observed GWAS results in UK Biobank. FMR was applied to summary statistics from the interim release (N=145k) in order to predict the results of the full release (N=460k). (a) Proportion of heritability explained by genome-wide significant SNPs 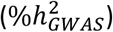. (b) Number of genome-wide significant loci (*M_GWAS_*). Error bars indicate 95% confidence intervals. For numerical values, see Table S1.

Predictions of 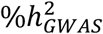 depend most strongly on the estimated distribution of effect sizes near the significance threshold, so we sought to determine whether it accurately estimates other portions of the effect size distribution. To evaluate its estimates of the tail of the distribution, corresponding to large-effect SNPs, we predicted 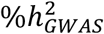 and *M_GWAS_* at more-stringent significance thresholds (*χ*^2^ > 100, 300, 1000). These predictions remained highly accurate (Figures S3-S4), indicating that FMR accurately estimates the tail of the HDM. We could not perform the same analysis for less-stringent significance thresholds because the observed values (computed using pruning + thresholding; see Methods) would be seriously biased due to false positives and winner’s curse^25,33^. We were most concerned that FMR estimates of the bulk of the HDM would be biased in a power-dependent manner, as there is no power to detect individual SNPs with effect sizes smaller than ~1/*N*. (However, power-dependent bias at small effect sizes was not observed in simulations, e.g. in Figure 1f). We expected that power-dependent bias, if it were a problem, would be stronger for FMR estimates derived from the N=145k vs. N=460k summary statistics, so we compared FMR predictions at N=145k vs N=460k of what 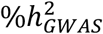 would be at very large sample size (N=2-10M). The predictions were nearly identical (Figure S5). These results indicate that FMR produces robust estimates across the entire range of the effect-size distribution.

We compared FMR predictions with the state-of-the-art method, GENESIS^14^, which models the distribution of causal effect sizes across SNPs using a 2- or 3-component Gaussian mixture model. For most traits, the GENESIS 3-component model is more realistic than the commonly used point-Normal model^13,17,18,20^, which assumes that all non-null SNPs have similar effect sizes. We applied GENESIS with the 3-component model to the interim-release UK Biobank summary statistics. For 6/22 traits, GENESIS threw errors after producing invalid (non-positive semidefinite) parameter covariance estimates. Across the remaining 16 traits, predictions of *M_GWAS_* and 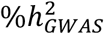 were upwardly biased by about a third (30% for *M_GWAS_* and 36% for 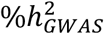), while FMR was approximately unbiased (2% and 7%) (Figure S6 and Table S1). An additional advantage of FMR is its computational efficiency; it runs in less than a minute, while GENESIS requires more than a day.

Uncorrected population stratification and cryptic relatedness can potentially confound methods such as FMR that estimate polygenic architecture. Similar to LDSC^26^, FMR models these confounders as excess noise in the effect-size estimates, and it assumes that this noise is uncorrelated with LD scores^26^. Unlike LDSC, FMR additionally assumes that stratification-induced effects follow a normal distribution, which is expected in the absence of subpopulation-specific selection^34^. In order to test these assumptions, we applied FMR to summary statistics for height from the GIANT consortium^2^ (2010 release, N=131k). Unlike the UK Biobank summary statistics (computed using BOLT-LMM^35,36^), these data contain strong, uncorrected population stratification that has led to confounding in studies of polygenic selection^37,38^. However, FMR produced nearly identical estimates on the two datasets (Figure S7), indicating that it is not confounded by population stratification within European populations.

A simplified version of FMR, FMR-noLD, can be used to estimate the distribution of marginal effect sizes across SNPs (see Methods). This method can be used to predict the QQ plot that will be observed at larger sample size. We applied FMR-noLD to interim-release UK Biobank summary statistics and predicted the QQ plots that would be observed at N=145k and N=460k. Predictions closely matched the observed QQ plots (Figure S8).

## Sample size targets across 37 diseases and complex traits

We applied FMR to publicly available summary statistics for 31 diseases and complex traits, including 21 UK Biobank traits (N=460k) and 10 European-ancestry meta-analyses (Table S3). We confirmed that FMR accurately estimated 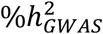 and *M_GWAS_* at current sample size for these traits (Figure S9-S10). We predicted the effective sample size required for genome-wide significant SNPs to explain 50% of heritability (Figure 3a, Table S5). This number varies by two orders of magnitude across traits, from a few hundred thousand to over ten million. Assuming a large number of controls, the number of cases required to explain 50% of disease heritability (i.e. *N_eff_*/4) was approximately one hundred thousand for IBD and hypothyroidism (which are highly heritable autoimmune diseases), several hundred thousand for schizophrenia and CAD (which have greater polygenicity), and several million for Alzheimer’s disease (which has low heritability in our dataset). We also estimated the sample size required for 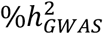 to reach 90% and determined that this number was ~10 *×* higher (Figure 3a), as a substantial fraction of heritability is explained by SNPs with extremely small effect sizes (see below). This difference implies that GWAS will have diminishing returns (in 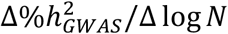) well before 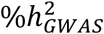 nears 100%, and it may be unnecessary to pursue 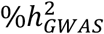 beyond 50% (see Discussion).

**Figure 3:**
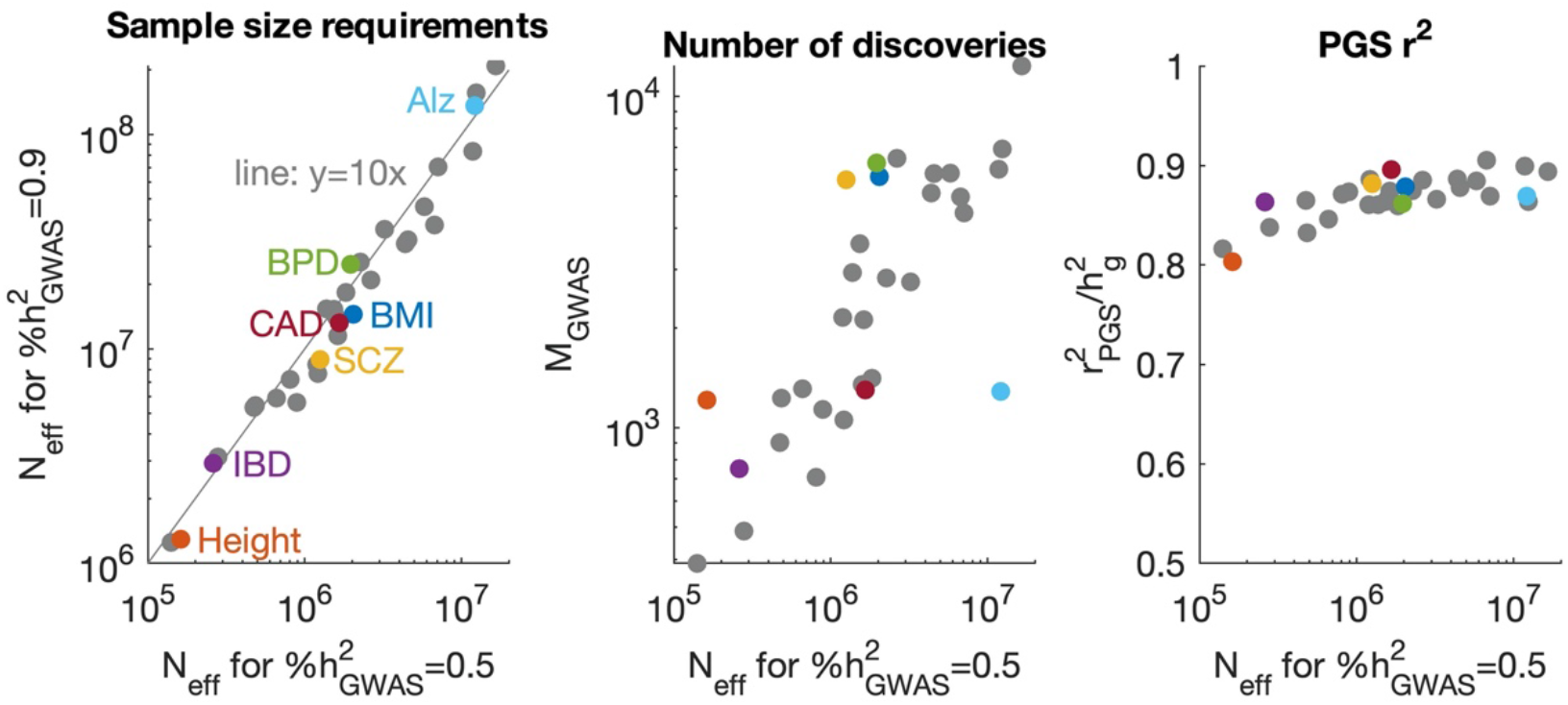
Sample size requirements and predictions for future GWAS. (a) Sample size required for genome-wide significant SNPs to explain 50% or 90% of SNP-heritability. (b) Predicted number of discoveries at the sample size where 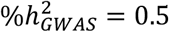. (c) Theoretical maximum PGS accuracy, defined as 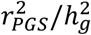, at the sample size where 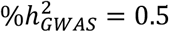. Existing PGS methods may perform less well than the maximum. For diseases, *N_eff_* is twice the harmonic mean of *N_case_* and *N_control_*, and 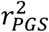 is defined on a liability scale. Numerical results are presented in Table S6.

We estimated the number of associated loci that would be discovered at the sample size when 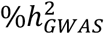 reaches 50%. This number ranges from a few hundred to several thousand for most traits (Figure 3b). The number of loci that would be discovered when 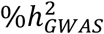 reaches 90% is roughly an order of magnitude larger (Table S6), although we caution that at such large sample size, independent associations would frequently be in partial LD, making them difficult to count. We applied FMR-noLD to estimate the number of SNPs (as opposed to loci) that would be genome-wide significant at very large sample sizes. When 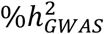 is 50%, the fraction of significant common SNPs will be 0.01-0.15 for most traits (Figure S11a), increasing to 0.1-0.5 when 90% of heritability is explained (Figure S11b). When ^~^10% of SNPs are genome-wide significant, distinctions between loci based on effect size may be more informative than those based on the significance threshold.

We derived an upper bound for the prediction r^2^ of any polygenic score (PGS) as a function of the HDM (see Methods). This bound is expected to accurately predict the within-population performance of an optimal PGS method in large samples, although existing PGS methods (e.g. LDpred^21^, which uses a point-normal model) may not approach this bound. We found that when 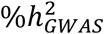 reaches 0.5, an optimal PGS would explain 80-90% of SNP-heritability for most traits (Figure 3c). Therefore, a 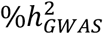 target of 0.5 should suffice to nearly maximize within-population PGS accuracy, assuming that optimal methods can be developed. Four of the UK Biobank traits we analyzed already have 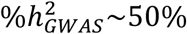, and a previous study^35^ reported the PGS accuracy of BOLT-LMM^36^ trained on these traits (BOLT-LMM improves association power by conditioning on a PGS that assumes a point-normal model). Reported r^2^ values were smaller than the predicted optimum (Table S7). This difference indicates that existing PGS methods have substantial room for improvement, possibly because the point-normal model is a poor approximation for real effect-size distributions (see below).

Predicted sample size requirements are subject to two notable sources of uncertainty. First, heritability may differ across different studies of the same disease, for example due to different age distributions of the participants, and mixed models also lead to increased effective heritability^35,36,39^. Increasing the heritability has the same effect on power as increasing the sample size. We report requirements for 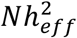, the product of effective sample size and heritability, in Table S5. Second, uncorrected population stratification and cryptic relatedness increase 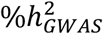, and recalibration using genomic control^40^ or the LDSC intercept^26^ decreases it. Our projections assume that the amount of uncorrected population stratification – in particular, the LDSC intercept minus one – will remain constant at increasing sample size, and that there will be no post hoc recalibration. If population stratification is poorly controlled and there is no recalibration, then 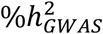 may appear to increase more quickly than expected, but some of the discoveries would be false positives. Conversely, if summary statistics are recalibrated to avoid false positives, then 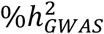 may fail to reach desired targets, as SNPs with very small true associations will never be distinguishable from those that are associated due to stratification. Therefore, it is crucial that future studies control confounding directly, through the use of mixed models^39^, principal components computed in the entire study population^41^, or family-based designs^42^.

## True-positive rates of non-significant loci

Genome-wide significance is a conservative threshold, and sub-GWS loci may represent true associations^1^. To quantify true positive rates at different significance thresholds, we define the *not-by-chance true positive rate* (NTPR): out of all loci exceeding a threshold, the proportion that are true positives with a correctly-estimated direction of effect, not counting “true-by-chance positives” whose estimated direction could just as well have been the opposite. The NTPR is related to the false sign rate^30^, and it can be estimated using FMR (see Methods for a mathematical definition and details about estimation).

The NTPR of nominally significant loci (*χ*^2^ > 4, roughly *p* < 0.05) would be nearly zero in an underpowered GWAS, as thousands of false positives are expected. However, current GWAS are powered to detect a vast number of true positives at this threshold; for height and BMI, they outnumber false positives (and true-by-chance positives), and the NTPR is >50% (Figure 4a). For most other traits, the NTPR of nominally significant loci was between 0.1 and 0.5. The proportion of heritability explained by these SNPs was only 57% on average; even in fairly well-powered studies, around 43% of heritability is explained by SNPs with effect sizes are so small that they are not even nominally significant. These estimates indicate that as GWAS sample size increases, multiple-hypothesis testing becomes a less serious problem. They underscore the tension between large sample size (which allows GWAS to detect many SNPs with small effect sizes) and extreme polygenicity (which causes missing heritability to persist despite ever-larger studies).

**Figure 4:**
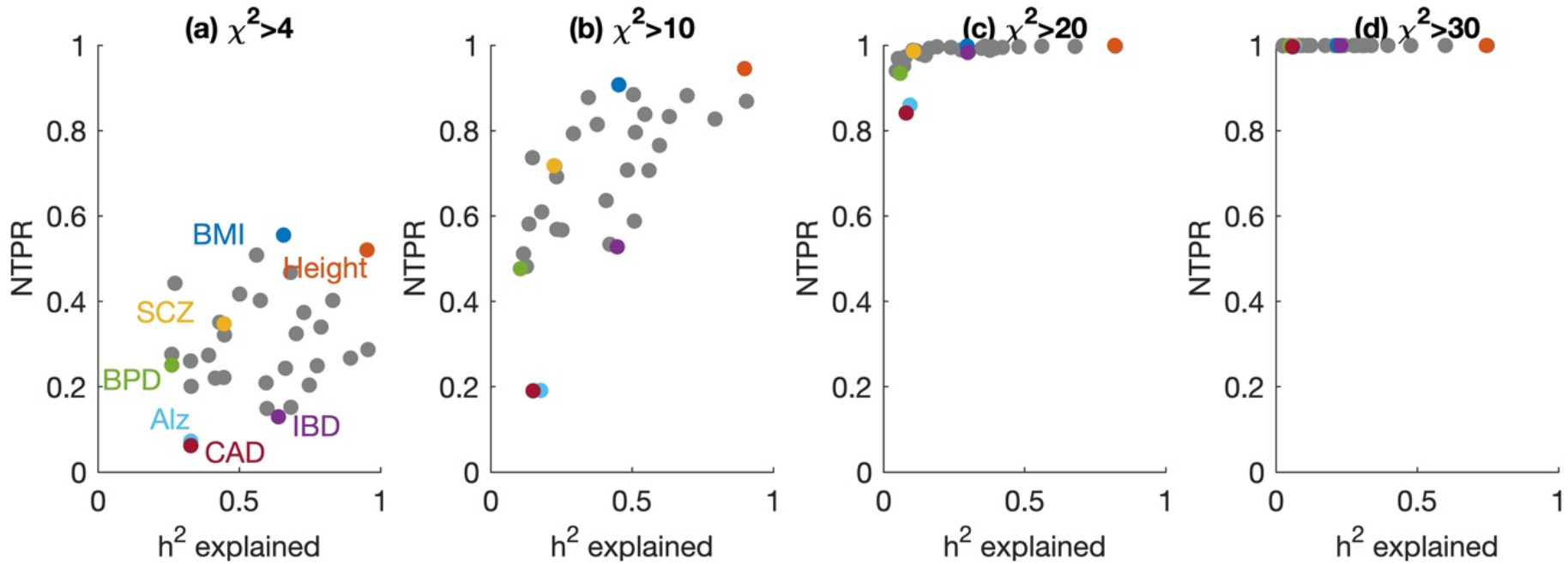
True-positive rates of loci at different significance thresholds. We quantify the heritability explained and the not-by-chance true positive rate for loci that exceed four significance thresholds, from nominal significance (a) to genome-wide significance (d). Numerical results are presented in Table S8.

We estimated the NTPR at larger significance thresholds. For most traits, it was greater than 50% at a *χ*^2^ threshold of 10 (Figure 4b) and greater than 95% at a threshold of 20 (Figure 4c). As expected, it was close to 100% for genome-wide significant SNPs (*χ*^2^ > 30) (Figure 4d). As sample sizes increase, genome-wide significance becomes increasingly conservative, and it may be permissible to relax the significance threshold as a function of sample size. However, population stratification and cryptic relatedness are more likely to generate false positives at increasing sample size, and it would be inadvisable to relax the significance threshold if confounding is poorly controlled.

We calculated the significance threshold that corresponds to an NTPR of 99%, assuming that confounders are well controlled. Across most traits, this number varies from 14 to 28, and SNPs exceeding this threshold explain 10-50% more heritability than those with *χ*^2^ > 30, with a 30-200% increase in the number of loci (Figure S12a-b). The difference is most pronounced for well powered, extremely polygenic traits such as BMI. We calculated what these quantities will be at the sample size when genome-wide significant SNPs explain 50% of heritability. The 99% NTPR threshold will range from 10 to 20, and SNPs exceeding this threshold will explain 60-75% of heritability (Figure S12c).

## The genetic effect-size distribution

We investigated the distribution of heritability across SNPs with different effect sizes. First, we estimated the median of the distribution: the minimum number of loci required to explain 50% of heritability, and the marginal effect size of a SNP at the 50^th^ percentile. The requisite number of loci ranges from a few hundred to several thousand, and the median effect size varies inversely (across 25 well-powered traits; see Methods) (Figure 5a). Psychiatric and brain-related traits had the greatest polygenicity, consistent with previous estimates^15^. These estimates exclude the MHC region, which has a large effect on autoimmune and inflammatory diseases.

**Figure 5:**
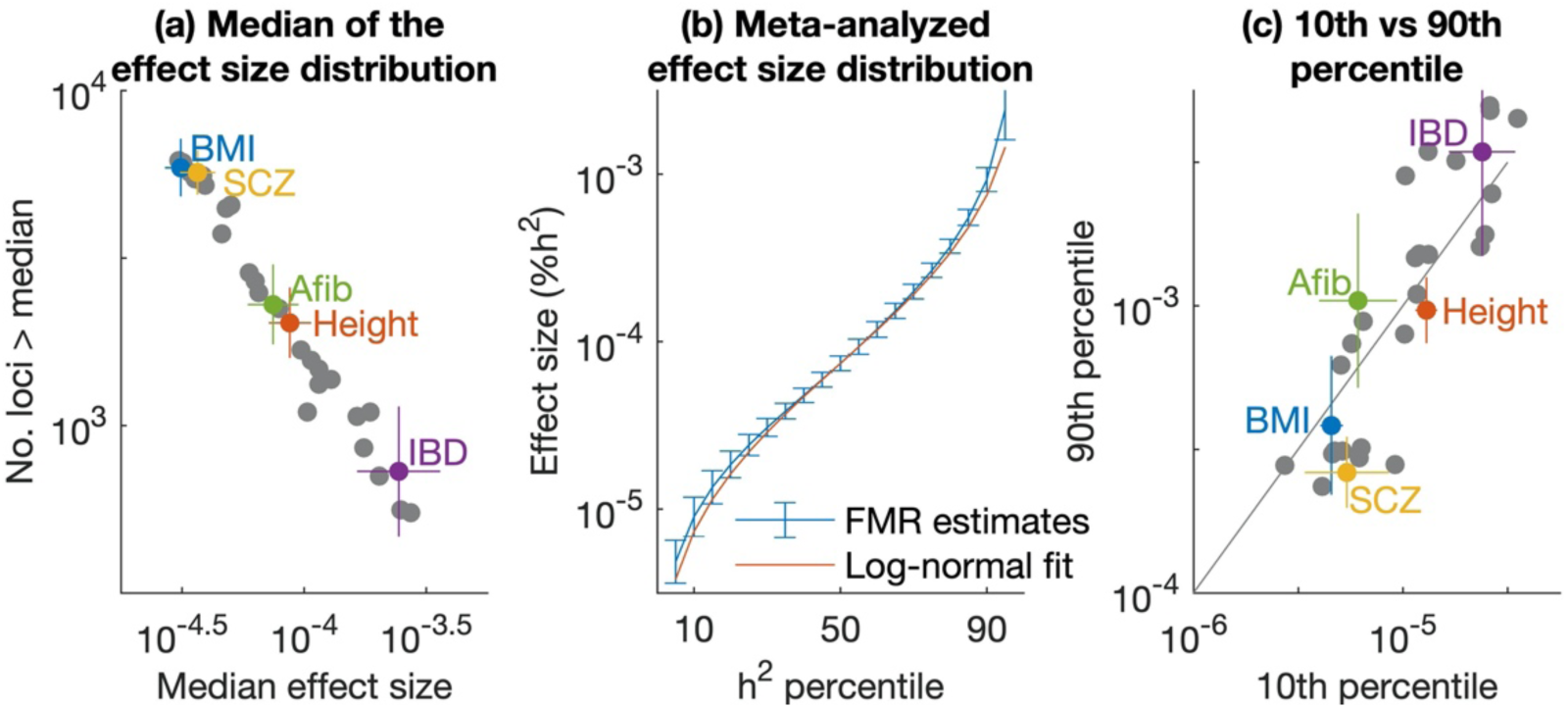
The shape of the heritability distribution. (a) Median of the effect size distribution: the effect size of a locus at the 50^th^ percentile of heritability, and the number of loci required to explain 50% of heritability. Estimates are shown for 25 well-powered traits (see Methods). Error bars indicate 95% confidence intervals, and they are only displayed for selected traits (see Table S10 for numerical results). (b) Quantiles of heritability, meta-analyzed across all 32 traits. We also show the quantiles that are expected under a log-normal model. (c) Effect sizes at the 10^th^ vs. 90^th^ percentiles of heritability for well-powered traits.

We asked whether heritability is concentrated within a narrow range of effect sizes, or whether it is dispersed across SNPs with a wide range of effect sizes. We computed meta-analyzed estimates of the effect-size distribution (across the 32 traits) (Figure 5b). On a log-10 scale, the 10^th^, 50^th^, and 90^th^ percentiles of heritability are −5.05 (s.e.=0.06), −4.13 (0.02), and −3.03 (0.04) respectively; that is, there is a factor-of-10 difference in effect size between the 10^th^ vs. 50^th^ and 50^th^ vs. 90^th^ percentiles, and a factor-of-100 difference between the 10^th^ vs. 90^th^ percentiles (95% CI: 80-130). These estimates are consistent with the factor-of-10 difference in sample size required to explain 90% vs. 50% of heritability (Figure 3a). The 5^th^ vs 95^th^ percentiles differ by a factor of ~500 (95% CI: 320-800). Thus, heritability is spread across SNPs with a wide range of effect sizes, and most heritability is explained by SNPs with effect sizes that are 1-2 orders of magnitude smaller than those discovered thus far.

We compared the range of the effect-size distribution across traits. The estimated difference between the 10^th^ vs 90^th^ percentiles of heritability was close to 100×, within a factor of ^~^2, for all well-powered traits (Figure 5c). The difference was slightly smaller for extremely polygenic traits than for those that are only moderately polygenic (*p* = 10^−4^; see Methods), possibly reflecting the finite number of loci in the genome.

We sought to identify a simple parametric model that fit the observed effect-size distribution, with simplifying assumptions about LD (see Methods). First, we considered the widely-used point-normal model^13,18,20,21,43^, even though it has been shown to fit poorly for many traits^14^. Under this model (with any parameter values), the 10^th^ vs 90^th^ percentiles differ by a factor of only 11, and the 5^th^ vs 95^th^ percentiles by a factor of just 22; it misspecifies the range of effect sizes by an entire order of magnitude.

Next, we considered a normal distribution over the log of *β*^2^, with mean *μ* and variance *σ*^2^. Under this distribution, every SNP has a nonzero effect size, but large values of *σ*^2^ lead to approximately-sparse distributions, because the range of effect sizes spans more orders of magnitude. Conveniently, if *β*^2^ follows a log-normal distribution over SNPs, then it also follows a log-normal distribution over components of heritability, with mean *μ* + *σ*^2^ and variance *σ*^2^ (see Methods). We fit a log-normal approximation to the FMR estimates, with a trait-specific mean and a constant variance parameter (see Methods). We found that the model provided an excellent fit to most of the distribution, both in a meta-analysis (Figure 5b) and for individual traits (Figure S13). It fit less well in the right tail of the distribution, underestimating effect sizes in the top 5% of heritability (Figure S13d). Nonetheless, the log-normal model is an attractive alternative to the widely-used point-normal model. It has the same number of free parameters (if its variance is fixed), it provides a far better fit to the true effect-size distribution, and it allows a straightforward conversion between distributions over SNPs vs. over components of heritability.

## Discussion

Our estimates suggest plausible sample-size targets for future GWAS. In order for genome-wide significant SNPs to explain 50% of disease heritability in populations of European ancestry, hundreds of thousands of cases will be required for most diseases (Figure 3a). Such numbers seem attainable. When they are achieved, thousands of additional loci below the genome-wide significance threshold will be probable true positives (Figure S12), and polygenic risk scores will explain up to 90% of heritable risk (Figure 3c). On the other hand, millions of samples would be required in order to increase 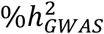 to 90% (Figure 3a), and for a disease at ^~^1% prevalence, this number could exceed the number of cases in Europe. Significant SNPs would span much of the genome (Figures 3b and S11), limiting their value. Extremely large sample sizes may be more useful in sequencing studies^44^ and in studies of diverse populations^45–47^.

Methods for polygenic risk prediction^6,21,48–50^ and fine mapping^7–9,25,27,28,51–53^ rely on models for the effect-size distribution. Several of them rely on a point-normal model: LDpred^21^ assumes this model for the purpose of polygenic prediction, and state-of-the-art fine mapping methods^9,51–53^ assume similar models with additional constraints. An alternative prior is the log-normal distribution, which fits the effect-size distribution well (Figures 5b and S13). Fine-mapping methods rely on a distinction between causal and non-causal SNPs both to estimate the posterior local effect-size distribution and to define interpretable estimands, such as minimum credible sets^9,27,28^. However, the binary distinction between causal and non-causal SNPs will become less tenable as sample sizes increase: as the number of associated loci increases exponentially, so too will the number of detectable causal SNPs per locus^27,28^, and it would be preferable to avoid relying on sparse priors, especially for the purpose of definition. For example, a non-sparse alternative to the minimum credible set is the minimum set of SNPs whose posterior-mean causal heritability is at least 95% of the total local heritability.

The range of effect sizes that explain 80% of heritability is approximately two orders of magnitude for all traits that we analyzed (Figure 5c), suggesting a shared underlying phenomenon. If heritability is mediated by cellular networks, as hypothesized by Boyle et al.^54^, then the size of the range may be determined by network architecture: suppose that the effect size of a gene decays exponentially with its network distance *d* from a set of disease pathways (or “core genes”^54^). If the number of genes at distance *d* increases exponentially with *d* (up to a point), then it could explain why heritability is distributed across such a large range of effect sizes, as the total heritability of genes at distance *d* is equal to their average effect size times the number of genes. The size of the range of effect sizes would be determined by network connectivity, i.e. the distribution of distances and the rate that effect sizes decay, which may be similar across trait-relevant cell types and subnetworks.

In recent work^15^, we showed that genetic effect-size distributions are “flattened” by negative selection, which prevents complex-trait heritability from being dominated by large-effect loci. Flattening could result either from direct selection acting on a trait (with selection coefficients approximately proportional to effect sizes) or from highly pleiotropic selection (with relative selection coefficients that vary widely). Direct selection is expected to produce a genetic architecture where most heritability is explained by SNPs with similar per-s.d. effect sizes, as the allele frequency of associated SNPs can rise until their per-s.d. selection coefficient is large compared with the rate of drift. On the contrary, we observed that heritability is spread across a wide range of effect sizes (Figure 5), more consistent with a model of primarily pleiotropic selection. Simons et al.^55^ also provided evidence that highly pleiotropic selection shapes complex-trait genetic architecture. It is possible for pleiotropic selection to strongly affect the genetic architecture of a trait even if that trait is also under direct selection.

Our study has several limitations. First, the HDM is defined as a distribution of marginal, rather than causal, effect sizes. This choice is desirable for projecting the results of future GWAS, and it simplifies the estimation task; however, inferences about causal effect-size distributions are also useful, for example to make inferences about cross-population genetic architecture^46,56^. We do not believe that any straightforward modification to FMR would allow it to estimate causal effect-size distributions. Second, FMR currently does not model SNP annotations, and in particular, it does not model LD-dependent architecture^57^, which can potentially bias heritability-style analyses^58,59^. Our real-data analyses (Figure 2) suggest that FMR produces reliable projections, and we previously found that estimates of *M_e_* (inversely proportional to the mean of the HDM) were extremely similar when we did not model LD-dependent architecture^15^. In principle, FMR could be extended to model any number of annotations, similar to S-LDSC^10^. Third, we have not made inferences about the joint distribution of effect sizes and allele frequencies. We estimate the distribution of per-s.d. effect sizes, and this is a natural choice because squared per-s.d. effect sizes are equal to per-SNP heritability, which determines association power; however, estimates of the joint distribution of effect sizes and allele frequencies are useful for making inferences about negative selection^15,18^. A stratified version of FMR would facilitate such inferences. Fourth, FMR requires fairly large GWAS sample size, so it is not applicable to small pilot studies of understudied diseases or populations. Despite these limitations, this study advances our understanding of genetic architecture, and it provides a glimpse of what will be found in future GWAS.

## Acknowledgements

I am grateful to Jenna Ballard, Ajay Nadig, Dan Weiner, Omer Weissbrod, Gaddy Getz, Ben Neale, and Eric Lander for suggestions and helpful discussions.

## URLs

Open-source software will be made available at github.com/lukejoconnor. GWAS summary statistics are available at alkesgroup.broadinstitute.org. GENESIS: github.com/yandorazhang/GENESIS.

## Methods

### Definition of the HDM

The HDM is formally defined under a non-i.i.d. normal model^15^ where per-s.d. causal effect sizes, denoted ***β***, follow a multivariate normal distribution:

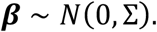

When Σ is a diagonal matrix, its diagonal entries, 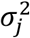, are equal to the expected heritability explained by each SNP. Per-s.d. marginal effect sizes are ***α*** = *R**β***, where *R* is the LD matrix. The non-i.i.d. normal model induces a heritability distribution over SNPs: SNP *j* is chosen with probability *p_j_* proportional to 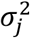. Let *J* be the index of the chosen SNP; the HDM is defined as the distribution of 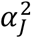. We write probabilities and expectations over this distribution using the notation *P*_*h*^2^_(·) and *E*_*h*^2^_(·) respectively.

The non-i.i.d. normal model makes essentially no assumptions about genetic architecture, as each SNP has a fixed variance parameter (instead of a fixed effect size), and arbitrary covariance is even permitted. The HDM is modeled as a finite mixture distribution for the purpose of estimation (see below), but no such assumption is needed for the purpose of definition. Under this model, the fixed parameters *R* and Σ have a certain symmetry: let *S* = *R*Σ. The heritability is *h*^2^(*R*, Σ) = *Tr*(*S*) = *h*^2^(Σ, *R*). The effective number of independently associated SNPs, a metric of polygenicity that we defined previously^15^, is 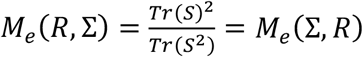. *M_e_* is equal to the SNP-heritability divided by the mean of the HDM, *E*_*h*^2^_(*α*^2^).

50% of random-effect SNP-heritability (*h*^2^) is explained by SNPs with marginal effect sizes less than the median of the HDM:

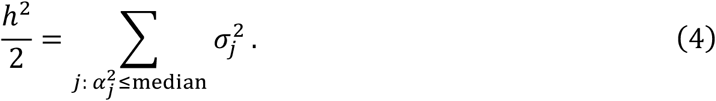

More generally, the cumulative distribution function (CDF) of the HDM is:

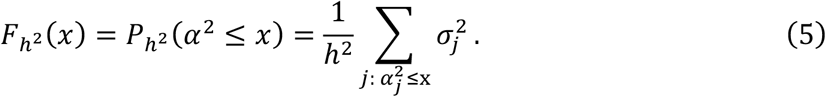

The non-i.i.d. normal model can allow for arbitrary covariance among SNP effect sizes. Let *S* = *R*Σ. When both *R* and Σ have nonzero off-diagonal entries, the expected heritability of SNP *j* is not 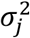, but rather the *j*th diagonal element of *S*:

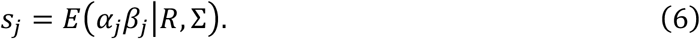

*s_j_* can be negative when SNPs with positively-correlated effect sizes are in negative LD, or vice versa, so it no longer defines a probability distribution. However, we can still define *E*_*h*^2^_ (η) as a weighted average (with possibly-negative weights) and *P*_*h*^2^_(·) as a weighted average of the appropriate indicator function:

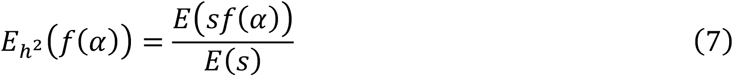

where *E*(·) denotes a uniform average across SNPs.

We define the heritability characteristic function:

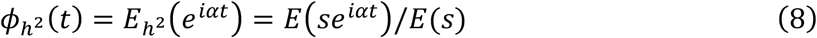

where *t* is a “sampling time.” *ϕ*_*h*^2^_ provides an equivalent way to describe the HDM as the cumulative distribution function, and it is useful for estimation (see below).

### Fixed vs. random-effect components of heritability

The HDM directly describes proportions of random-effect heritability explained by SNPs with various effect sizes. For example, the random-effect heritability explained by genome-wide significant SNPs is:

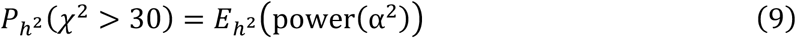

where “power” is the GWAS power function at a given sample size, and *E*_*h*_^2^ denotes an expectation over the HDM (i.e. a weighted average, with weights proportional to 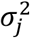).

However, we distinguish between random-effect components of heritability (sums of *σ*^2^) and fixed-effect components of heritability (sums of *β*^2^, or more precisely sums of *αβ*). For example, the fixed-effect heritability of GWS SNPs is generally larger than the random-effect heritability, and when we refer to 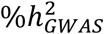, we mean the fixed-effect heritability.

In detail, the random-effect proportion of heritability of a set of SNPs *A* is:

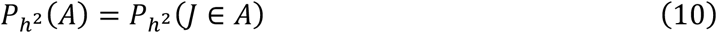

where *J* is the random index of a SNP chosen with probability proportional to *s*. The fixed-effect proportion of heritability is:

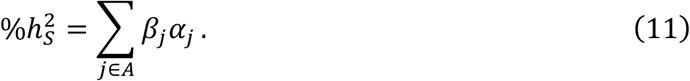

If *A* is fixed, *P*_*h*^2^_(*A*) is a fixed constant equal to 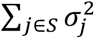, and 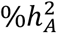 is a random variable whose expectation is equal to *P*_*h*^2^_(*A*). However, *A* itself might itself be random (e.g. the set of SNPs that are genome-wide significant in a study). In that case, *P*_*h*_^2^(*A*) is also a random variable, and it is generally false that *E*(*P*_*h*^2^_(*A*)) = *E*(%*h*^2^(*A*)). For example, if *S* is the set of genome-wide significant SNPs, then *E*(*P*_*h*^2^_(*A*)) = *E*(%*h*^2^(*A*)), as SNPs that reach genome-wide significance are likely to have larger effect sizes than those that do not even conditional on *σ*^2^.

Similarly, we choose to describe the effect-size distribution using quantiles of fixed-effect heritability, and these quantiles differ from those of the HDM. We make this choice because fixed-effect heritability is what actually exists in the population, and it is not as useful to quantify proportions of heritability that might be expected in a theoretical population whose effect-sizes were resampled from the same distribution. The quantiles of the HDM are generally smaller than those of fixed-effect heritability, especially in the left tail (corresponding to small-effect SNPs). For the heritability explained by genome-wide significant SNPs, there is an additional rationale for measuring fixed-effect heritability: it is actually possible to estimate this quantity directly (not using FMR) via pruning and thresholding, and for a quantity whose purpose is to benchmark progress, this property is essential.

### Derivation of estimation equation

We derive equation (3) and discuss the assumptions that are required. Let ***r*** denote a column of *R* chosen uniformly at random, and let ***r***^2^ denote the element-wise squared column vector. Under an LD approximation (see below), the following holds:

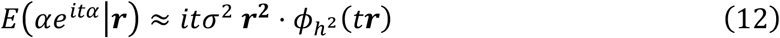

where *σ*^2^ = *h*^2^/*m*.

To derive this, we evaluate the derivative of *E*(*e^iαt^*|***r***) in two ways. First:

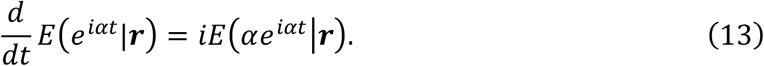

Second, we substitute the normal characteristic function before taking the derivative. Let 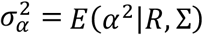 then 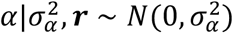, and:

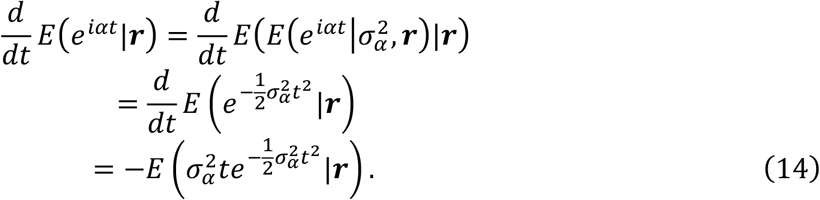

Now we use the LD approximation: suppose that LD between causal SNPs is either strong or weak, such that for our randomly-chosen SNP and all other SNPs *j*, either *S_j_r_j_* ≈ 0, or *α* ≈ *r_j_α_j_*. This type of approximation – which holds under two extremes, corresponding to weak LD or strong LD between causal SNPs – is expected to work well in practice because there is a limit to how strongly it can be violated. A similar assumption is used for LD 4^th^ moments regression, and it was found to be robust^15^. The LD approximation is used as follows:

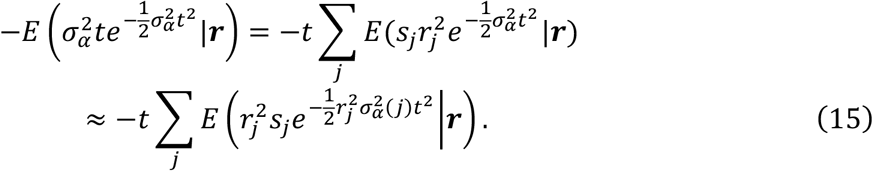

We make the further “representative-SNPs” assumption, that within the summation, *E*(·*_j_* |***r***) = *E*(*·_j_*) (i.e., SNPs tagged by regression SNPs are representative of all reference SNPs). A similar assumption is used by LD score regression and similar methods^10,15,26^, but it can be violated in the presence of LD-dependent architecture^57,60^, because SNPs with high LD are more likely to be tagged by regression SNPs, which leads to bias if their effect-size distributions are non-representative of the effect size distribution of all reference SNPs. In principle, this assumption could be improved by stratifying the analysis across LD-related annotations in the Baseline-LD model^57^, but FMR currently does not allow stratified analyses. We previously found that LD4M (which estimates the mean of the HDM) produces extremely similar genome-wide estimates when LD-related annotations are included or excluded^15^. It may be more critical to model LD-dependent architecture when performing stratified analyses of LD-correlated functional annotations such as DHS^26,57,61^.

Under the representative-SNPs assumption, we have:

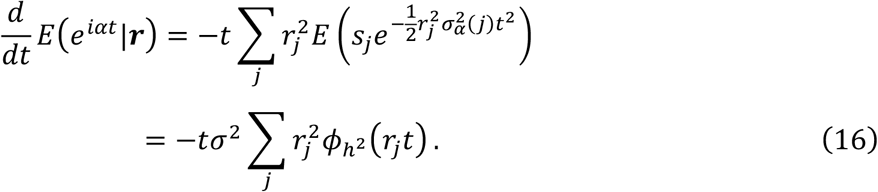

Now, suppose that *f*_*h*^2^_ is a mixture distribution with unknown weights:

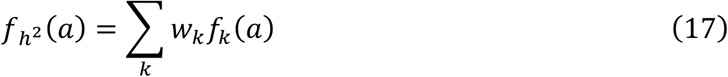

where *f_k_* denotes the PDF of the *k*th mixture component, and *w_k_* denotes its mixing weight. Then equation (3) takes the form

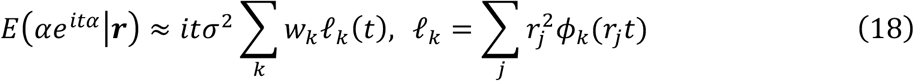

where *ϕ_k_*(*t*) is the characteristic function of the kth mixing component (e.g. 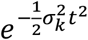 for a normal distribution). This equation implies that if *α* were given, we could estimate *w* by regressing the LHS on the LD scores for some appropriate values of *t*.

At finite GWAS sample size, we do not observe ***α***, but rather a noisy estimate:

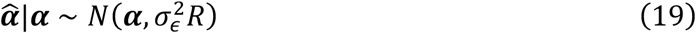

where 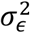 is 1 divided by the GWAS sample size in the ideal case (in practice, we estimate it using the LD score regression intercept^26^). Let 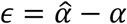. We account for sampling noise by observing that:

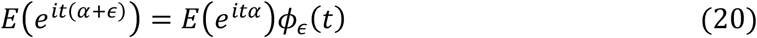

where 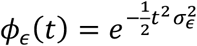. Therefore, if

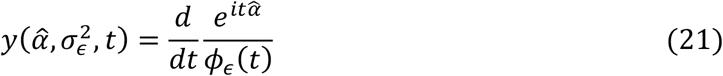

then

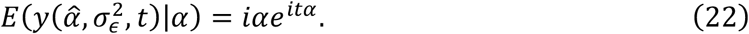

We obtain the estimation equation:

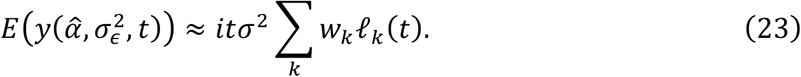

### Finite reference panel size

In practice, Fourier LD scores are estimated from a reference panel of finite size, so the in-sample LD matrix differs from the population LD matrix. We wish to obtain an approximately unbiased estimate of the population Fourier LD scores.

For a normal mixture component with variance *σ*^2^, the Fourier LD score is defined as:

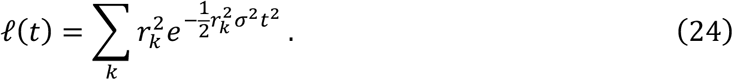

The sample correlation, 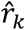, approximately follows a normal distribution with mean *r_k_* and variance 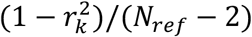, where *N_ref_* is the size of the reference panel. Define 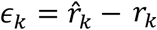. An unbiased estimate of ℓ(*t*) can be obtained by taking the obvious biased estimate:

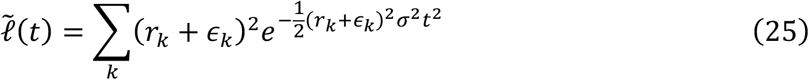

and expanding each term around *ϵ_k_* = 0:

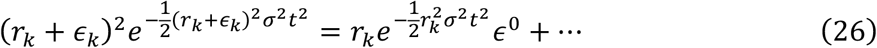

The term we must correct for is the *ϵ*^2^ term, whose coefficient is:

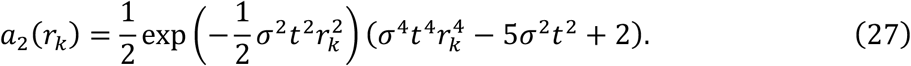

We obtain a bias-corrected estimate of ℓ:

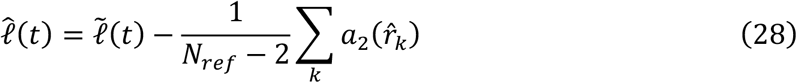

which is approximately unbiased:

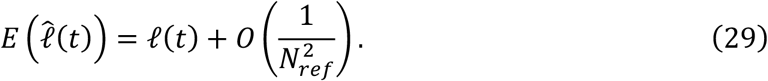

### Fourier mixture regression

In detail, FMR involves the following steps:

1. Choose *K*_1_ mixture components and *K*_2_ sampling times. We recommend using normal mixture components with *σ*^2^ values in a geometric series, and choosing one sampling time for each component as 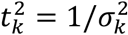. The scale of the *σ*^2^ values does not matter (except that very large or small values can lead to numerical issues), as the summary statistics will be scaled to match the *σ*^2^ values. The use of a finite number of sampling times is analogous to approximating a probability density function using a histogram with finitely many bins. The choice to use *σ*^2^ and *t*^2^ values in the same geometric series is convenient because it reduces the number of Fourier LD scores that must be computed: for the normal distribution characteristic function, *ϕ_k_*(*t*) depends only on 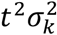, so many pairs 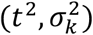 have the same value of ℓ*_k_*(*t*), and the number of nonidentical Fourier LD scores that must be computed is only *K*_1_ + *K*_2_ – 1 instead of *K*_1_*K*_2_. We use *K*_1_ = *K*_2_ = 13 and 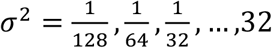. (The summary statistics are scaled so that the value “1” in this series corresponds to the mean of the HDM).
2. Compute Fourier LD scores from a reference panel. We use European-ancestry samples from the 1000 Genomes Project^31^. We do not impose any allele-frequency threshold in this step, but we do restrict the set of regression SNPs to common SNPs (MAF>0.05), which has the effect that our estimates describe the common effect-size distribution because there is little LD between common and rare SNPs. (We confirmed that nearly-identical estimates are obtained if a 1% minor allele frequency threshold is imposed on reference SNPs).
3. Use LD 4^th^ moments regression to estimate the mean of the HDM (we previously called this the “average unit of heritability”). Divide the summary statistics by this value. Compute the LD score regression slope, 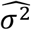, and intercept, 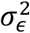, on the same scale.
4. For each SNP and each value of t, compute 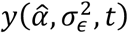 (equation 21). Concatenate into a vector *y* of length *MK*_2_.
5. Arrange the Fourier LD scores, ***ℓ**_k_*(*t*), into a matrix 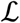 of size *MK*_2_ × *K*_1_. Each column corresponds to a mixture component, and row *m* + (*k* – 1)*M* corresponds to SNP *m* and sampling time *t_k_*.
6. Let ***ℓ***^(2)^**1**_1×*K*_1__ be the vector of LD scores tiled *K*_1_ times to obtain a matrix of size *M* × *K*_1_, and concatenate to the bottom of 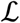. Concatenate 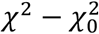 to the end of *y*. This corresponds to including the LD score regression moment equations 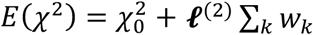.
7. Let 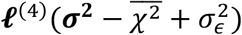 be the outer product of the vector of LD 4^th^ moments and the vector of variance parameters minus the mean *χ*^2^ statistic plus the LD score intercept, and concatenate to the bottom of 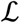. Concatenate 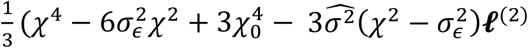 to the bottom of *y*. This corresponds to including the LD4M moment equations.
8. Set the regression weights. The weight of entry (*k* – 1)*M* + *m*, where *m* ≤ *M*, is proportional to 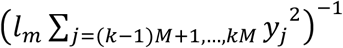, where *l_m_* is the in-sample LD score of SNP *m*. This heuristic weighting scheme down-weights SNPs with high LD (reducing redundancy at high-LD loci), and it weights the sampling times (as well as the LDSC/LD4M moment conditions) so that they are all about equally important in the objective function. An optimal weighting scheme would include off-diagonal weights accounting for correlations between sampling times, presumably resulting in reduced standard errors.
9. Let 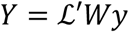 and 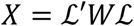, where 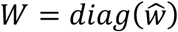. Use nonnegative weighted least squares to regress *Y* on *X*, which is equivalent to performing a weighted nonnegative regression of *y* on 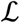; we perform this regression using the Matlab function *lsqlin*. The nonnegative regression does not increase the computational complexity, as the input to *lsqlin* has size *K*_1_ × *K*_1_ (whereas complexity is dominated by the factor of *M*). The HLA region is excluded from the regression.
10. To calculate standard errors, use a block jackknife with one contiguous block of SNPs left out for each jackknife iteration, as previously described^26^. We use 100 contiguous jackknife blocks with roughly an equal number of regression SNPs per block. Our implementation avoids recomputing any step whose complexity scales with *M*, such that the computational complexity does not scale with the number of jackknife blocks.

### Implementation of FMR

Open-source software for FMR is available (see Code Availability). FMR is implemented in MATLAB, and it requires the “lsqlin” function from the MATLAB optimization toolbox. The software does not require any installation, and scripts are provided to replicate the main results.

### Simulations

We perform simulations using real LD patterns computed from UK Biobank typed SNPs (M=455k) as previously described^15^. We simulated per-normalized-genotype causal effect sizes from:

1. An i.i.d. point-normal distribution with a specified proportion of causal SNPs (10%, 1% and 0.1% in Figure 1a-c). The variance parameter was equal to 0.1/*M_c_*, where *M_c_* is the expected number of non-null SNPs.
2. A four-component mixture of normal distributions with a null component and small-, medium- and large-effect non-null components (Figure 1d-f). In Figure 1d, the proportion of SNPs in each non-null component were 0.01, 0.01/5, 0.01/25 respectively. In Figure 1e: 0.01, 0.01/10, 0.01/100. In Figure 1f: 0.01, 0.01/50, 0.01/2500. The effect-size variance of each component was equal to 0.1/3*M_c_*, where *M_c_* is the expected number of SNPs in that component.

Then, we simulated GWAS summary statistics from the asymptotic sampling distribution^62^, which depends on the LD matrix, as previously described^15^. We calculated Fourier LD scores directly from the LD matrix that was used to generate the summary statistics, with no correction for finite reference panel size. We applied FMR with the same model and parameter settings as we used for analyses of real traits.

### Heritability explained by genome-wide significant SNPs

We compute the fixed-effect heritability explained by genome-wide significant SNPs in two ways. First, we perform significance thresholding followed by LD pruning. We compute a sparse LD-pruning matrix *Q* whose (i,j)th entry is equal to 1 if 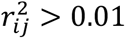 and zero otherwise. We sort all genome-wide significant SNPs by descending *χ*^2^ statistic. Iteratively, starting from SNP *i* at the top of the list, we discard all SNPs *j* from the remainder of the list such that *Q_ij_* = 1. After iterating to the bottom of the list, what remains is a pruned set of genome-wide significant SNPs. We estimate 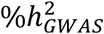 as the sum of their marginal effect-size estimates divided by the FMR/LDSC heritability estimate.

Thresholding and pruning is subject to three potential sources of bias. First, it is affected by winner’s curse due to thresholding: SNPs with true effect sizes near the significance threshold only pass the significance threshold when their effect size (and their heritability) is overestimated^33^. Second, it is affected by winner’s curse due to partial LD: the most significant SNP at a locus might differ from the causal SNP, leading to upward bias when a causal SNP is in strong-but-not-perfect LD with many tag SNPs. Third, as the number of significant loci becomes large, it is potentially affected by excessive LD pruning, which leads secondary signals in partial LD with a stronger signal to be discarded. This source of bias could be reduced by using a less-stringent LD threshold (we use 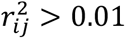), at the risk of introducing additional upward bias due to counting the same signal multiple times.

Second, we estimate 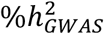 using FMR. Let power(*α*^2^, *N*) denote the GWAS power function for a SNP with effect size *α*^2^ and sample size *N*. We draw 100,000 samples *X* from a standard Normal distribution. For each heritability component *k*, with effect-size variance 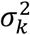, we compute the proportion of fixed-effect heritability that will be explained by genome-wide significant SNPs for that component:

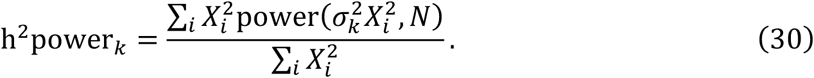

The FMR estimate of 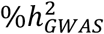 is

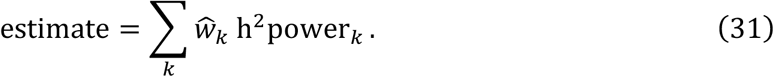

When forecasting the results of very large future GWAS, we assume that the LD score intercept will be used to calibrate the significance level and that it will remain constant from current sample sizes.

### The number of genome-wide significant loci

FMR estimates can be converted into an expected number of significant loci. If there is no LD, then the heritability density function, *f*_*h*^2^_(*x*), is proportional to the SNP-density function times *x*^2^:

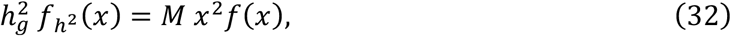

where 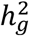 is the heritability and *M* is the number of SNPs. In the presence of LD, a similar equation does not generally hold, but we can interpret 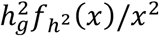 as the derivative of the number of loci. Define

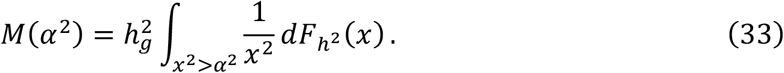

We interpret *M*(*α*^2^) as the expected number of independent associations with effect size greater than *α*^2^. Similarly, for a GWAS of sample size N, define

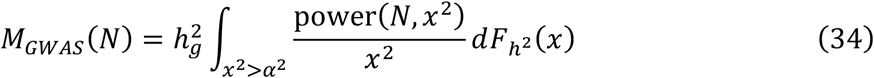

where power(*N*, *x*^2^) is the probability that a SNP with effect-size *x* will be genome-wide significant at a sample size *N*.

### FMR-noLD

A modified version of FMR can be used to estimate the distribution of marginal effect sizes. This estimation problem is straightforward, because it is unnecessary to account for LD. Although other methods are possible, we use a method-of-moments estimator similar to FMR. The estimation equation is:

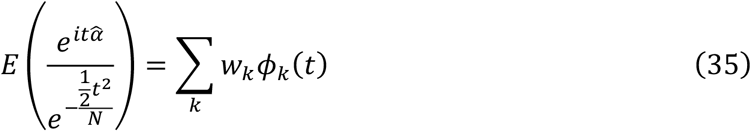

where *w_k_* is the mixing weight of component *k*. We compute the LHS for *K*_1_ sampling times and compute the mean *y* and covariance matrix *A* of the resulting *M* × *K*_1_ matrix. We compute a *K*_1_ × *K*_2_ matrix *X* whose entries are *ϕ*_*k*2_(*t*_*k*_1__). We calculate a weights matrix *W* = (*λI* + (1 – *λ*)*A*)^−1^, where *λ* = 0.1. We use nonnegative weighted least squares to regress *y* on *X* with weights *W*, producing an estimate of *w*. We use a mixture of normally distributed components with variance equal to 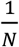 times 0,2^−7^, 2^−6^, …, 2^7^. We use sampling times equal to *N* times 2^−7^, 2^−6^, …, 2^8^. Reasonable choices of these parameters lead to very similar estimates of the distribution (although the specific estimate of *w* can vary widely).

### Risk prediction accuracy

The HDM allows us to calculate risk prediction accuracy and missing heritability as a function of GWAS sample size. If Σ is given, the optimal risk prediction method is straightforward: 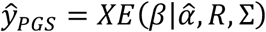. The risk prediction accuracy is:

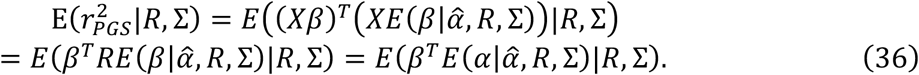

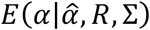 is the posterior of the mean of a multivariate normal random vector, 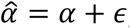, which itself follows a multivariate normal distribution with known variance 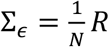.^62^ The formula for the posterior mean is:

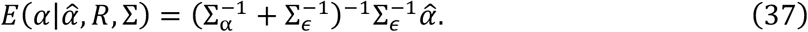

Assuming the various matrices are invertible, this becomes:

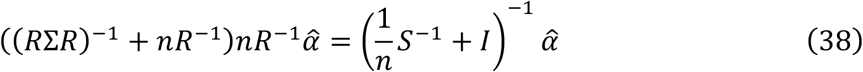

where *S* = Σ*R* = *E*(*βα^T^*). Therefore

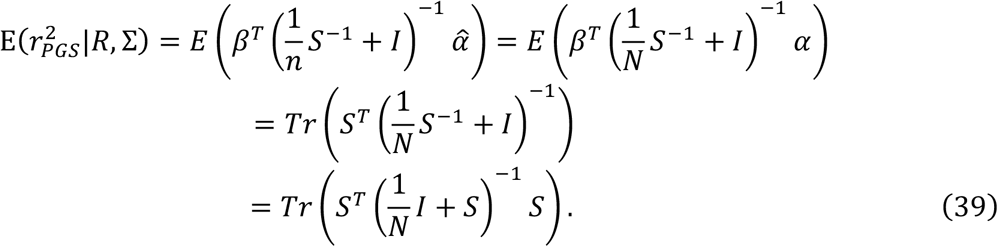

When *n* is small, we obtain an approximation that we derived previously^15^:

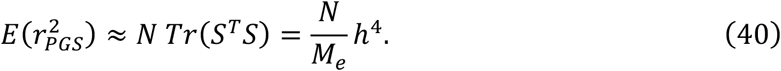

When *n* is large, we obtain a different approximation:

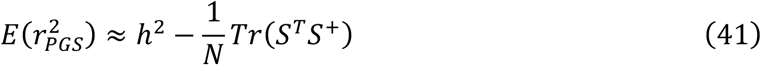

where we have replaced the inverse with the pseudo-inverse *S*^+^ to allow for non-invertible S. For example, under a point-normal model with *M_c_* causal SNPs and no LD, *Tr*(*S^T^S*^+^) = *M_c_*, and at large N the risk prediction accuracy is 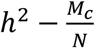.

Alternatively, we can use the FMR approximation, that LD between causal SNPs is strong or weak, which leads to the following approximation:

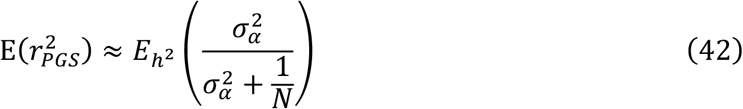

where 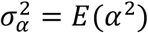, i.e. a diagonal element of *R*Σ*R*. Previously^15^, we gave the incorrect formula 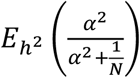. This formula is an approximate upper bound on the expected accuracy of any PGS method, and in practice, PGS methods are not given Σ, so their accuracy will be lower. We expect that the bound will be tight for large N and for the optimal method that is not given Σ, as the prior Σ becomes less important as more data is collected. However, existing methods may perform substantially less well if they badly misspecify the prior distribution.

### The not-by-chance true positive rate (NTPR)

The NTPR measures the proportion of two-tailed positive tests that correctly reject the null hypothesis in favor of the correct alternative, and not by chance. It is designed to satisfy the property that for any mixture of null and non-null effects, if there is no power to estimate the direction of effect, the NTPR is zero. It thereby avoids an arbitrary distinction between effects that are near zero 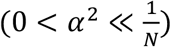 and those that are exactly zero. For a Z-score threshold *T* ≥ 0, NTPR is defined as the fraction of true positives with the correct sign *minus* the fraction of sign errors:

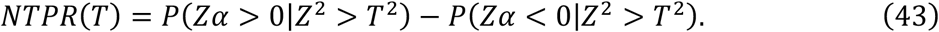

Under sign symmetry, for every near-zero effect whose effect direction is estimated incorrectly, there is expected to be a near-zero effect whose effect direction is estimated correctly due to chance, so subtracting the fraction of sign errors (counting them as −1 true positives, rather than as 0 true positives) cancels out the “true-by-chance” positives. False positives – SNPs whose true effect size is exactly zero - count as 0 true positives, so the expected number of true positives is the same for a SNP whose effect size is exactly zero or nearly zero.

The NTPR is related to the false sign rate (FSR) introduced by Stephens^30^. In this application, the FSR is:

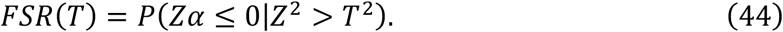

If there are no null effects (such that *P*(*α* = 0) = 0) then *NTPR* = 1 – 2*FSR*. However, we believe that the NTPR is a preferable definition, because it avoids an arbitrary distinction between zero and near-zero effects. If *α* is exactly zero for every SNP, then the NTPR is 0 and the FSR is 1. If *α* is extremely close to zero for every SNP, such that the sign of *Z* is independent of the sign of *α*, then the NTPR is still equal to zero but the FSR is equal to 1/2, because the direction of effect is correct by chance half of the time. In other words, the NTPR but not the FSR is a continuous function of the quantiles of *α*.

In practice, the importance of this distinction depends on whether the estimated distribution of *α* has a point mass at exactly zero. Our implementation of FMR-noLD does include a point mass at zero, but this could easily be modified to a normal distribution with variance extremely close to zero, and then the NTPR and FSR would be equivalent. Stephens observed that estimates of the FSR are more robust when all effects are assumed to be non-null, but argued that this assumption is anti-conservative^30^. This is the limitation that is addressed by the NTPR.

It is impossible to label each individual true positive as “by-chance” or as “not-by-chance,” but it is possible to define a local not-by-chance true positive rate (lntpr; analogous to the lfsr^30^), which is defined as

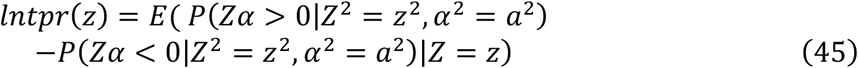

where the integral is computed over the posterior, *α*|*Z*. For an individual true positive, the lntpr measures the degree to which it is true by chance: it is zero if the sign of *z* was a coin flip, and one if the sign of *z* was guaranteed to equal the sign of *α* for any credible value of *α*. The NTPR is the expected value of the lntpr given *Z*^2^ > *T*^2^. The lntpr is related and the local false sign rate.

### Estimating the NTPR

The NTPR is estimated using FMR, with an additional assumption about LD structure. Suppose that the distribution of marginal effect sizes over SNPs is a mixture of normal distributions with variance parameters 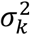 and weights *w_k_*. Let *X* ~ *N*(0,1). The NTPR over SNPs at Z-score threshold T and sample size N is:

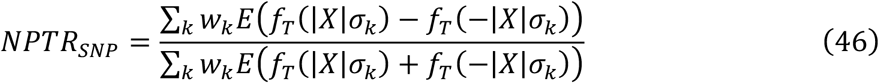

where 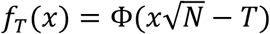 and Φ is the standard normal CDF.

*NPTR_SNP_* can be estimated using FMR-noLD, but it is not actually what we want: many nominally significant SNPs are in partial LD with a lead SNP that is much more significant, and these SNPs are much more likely to be true positives compared with a nominally significant lead SNP. Therefore, we define instead *NPTR_locus_*, which is defined under the simplifying assumption that the LD matrix is block-diagonal, with independent loci and perfect LD within each locus. This choice is necessary so that it makes sense to define a locus discretely; an alternate approach is to define *NTPR*_*h*^2^_ as an integral over the heritability distribution, but we believe that this definition would be difficult to digest.

Under the independent-loci approximation, the distribution of marginal effect sizes over loci is determined by the HDM: if the HDM is a mixture of normal distributions with variance parameters 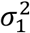 and weights *w_k_*, then the distribution over loci is a mixture of normals with variance parameters 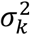 plus a point mass at zero, and the weights of the non-null components are proportional to 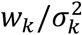. The weight of the null component is determined by the number of loci and the heritability, as the effect-size variances of the non-null loci sum to the heritability.

Real LD does not actually form discrete loci. In order to calculate the equivalent number of loci that should be used to calculate *NPTR_locus_*, we compute the number of SNPs divided by the harmonic mean of the LD score, or equivalently:

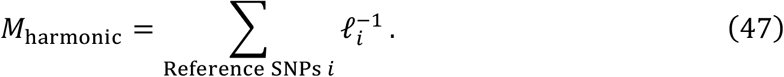

This definition has the property that if the genome does comprise independent loci with perfect LD, then *M*_harmonic_ is equal to the number of loci. It is equal to 4.0 × 10^5^ in our LD reference panel. Estimates of *PTR_locus_* are not highly sensitive to the specific value of *M*_harmonic_ (Table S8).

We compute *NPTR_locus_* by converting the FMR estimate of *w* and *σ*^2^ into an estimate of the distribution of effect sizes over loci, assuming that the number of loci is 4.0 × 10^5^, and evaluating equation (46).

### Well-powered traits

In Figure 5, we display results for well-powered traits whose 10^th^, 50^th^, and 90^th^ quantiles had standard errors less than 0.5 on a log-10 scale. We made this choice because estimates were very noisy for a few traits (especially at the 10^th^ percentile). Estimates for all traits are contained in Table S10.

### Meta-analysis across traits

We perform unweighted meta-analyses across traits on a log scale, i.e. by averaging the exponents. Standard errors of meta-analyzed quantities are estimated by jackknifing the mean; this approach has the advantage that it does not assume independent standard errors across traits. 95% confidence intervals are assumed to be +-1.96 standard deviations. When computing standard errors for a difference (e.g. the fold-difference between the 10^th^ and 90^th^ percentiles of heritability), we jackknife on the difference.

### Hypothesis testing

We performed a hypothesis test for the difference between the 10^th^ and 90^th^ percentiles of heritability comparing highly polygenic vs moderately polygenic traits. Highly polygenic traits were defined as well-powered traits whose estimated 90^th^ percentile was smaller than 10^−3.7^ (7 traits), and moderately polygenic traits were defined as the remaining well-powered traits (18 traits). This threshold was chosen after seeing the data. For the two sets of traits, we computed the difference between their 10^th^ and 90^th^ percentiles, took the difference between the differences, and estimated the standard deviation by jackknifing. We computed the p-value via a *χ*^2^ test.

### Properties of the point-normal model

We calculated the quantiles of heritability that would be observed under a point normal model by drawing 1 million samples from a *χ*^2^(1) distribution, sorting them, and computing the quantiles of their cumulative sum. For example, the 10^th^ percentile was equal to the *χ*^2^ value such that the sum of all smaller samples was 10% the total sum.

### The log-normal model

Suppose that there is no LD, and log*β*^2^ follows a normal distribution across SNPs. Letting *X* = log*β*^2^,

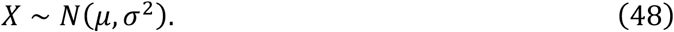

The PDF of *X* is:

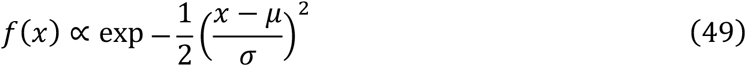

Let *g*(*x*) be the PDF of *X* over components of fixed-effect heritability; then

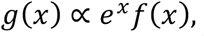

because the heritability explained by a SNP is equal to *β*^2^ = *e^x^*. It follows that

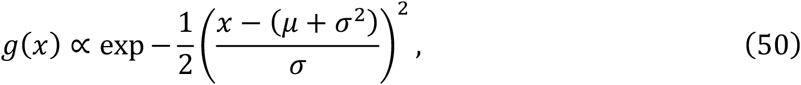

and that log*β*^2^ also follows a log-normal distribution over components of heritability, with its mean shifted from *μ* to *μ* + *σ*^2^.

In the presence of LD, and in particular LD between causal SNPs, there is no 1:1 correspondence between the distribution of causal effect sizes over SNPs and the distribution that we are able to estimate, i.e. the distribution of marginal effect sizes over components of heritability. A log-normal model may still be appropriate, but its parameters would have to be estimated using some other method.

To fit the log-normal model (over components of heritability), we match the 50^th^ percentile and the difference between the 90^th^ and 10^th^ percentiles. Each trait has trait-specific mean (i.e. *μ* + *σ*^2^) equal to the log of the FMR-estimated median. The standard deviation is fixed at 0.785 for every trait; this value is obtained by taking the log-scale difference between the 10^th^ and 90^th^ percentiles, meta-analyzed across traits, and dividing by 2.56, which is equal to the number of standard deviations between the 10^th^ and 90^th^ percentiles of the normal distribution.

