## Supplementary figures and tables for "The distribution of common-variant effect sizes"

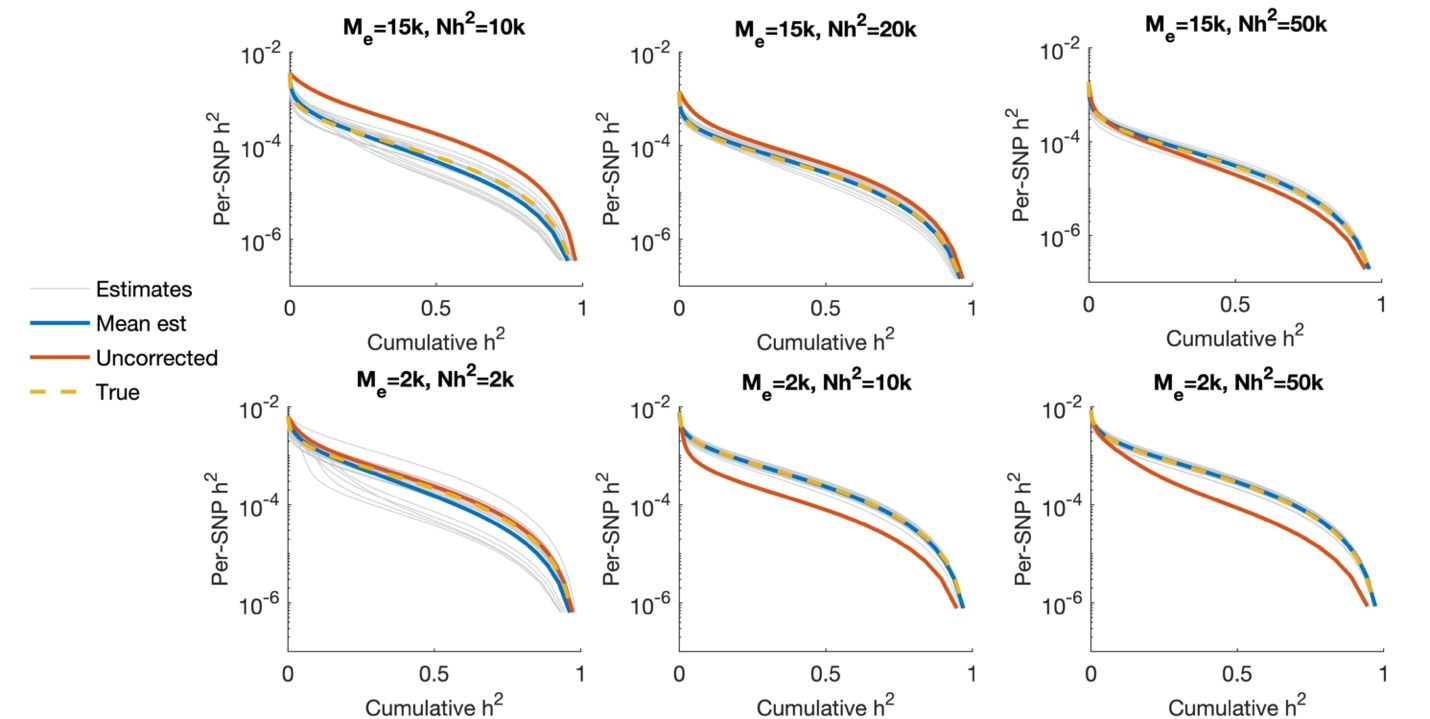

Figure S1: Performance of FMR in simulations at different sample sizes. We show the true HDM (yellow), estimates for 10 individual simulation replicates (grey), the mean estimate across 20 replicates (blue), and the mean uncorrected estimate, which was computed without any correction for finite sample size. Data were simulated under a point-normal model with either 1% or 10% of SNPs having nonzero causal effect sizes.

|  | GENESIS prediction (N=145k) | | FMR prediction (N=145k) | | Observed (N=460k) | |
| --- | --- | --- | --- | --- | --- | --- |
|  | Discoveries | %h2GWS | Discoveries | %h2GWS | Discoveries | %h2GWS |
| Eosinophils | 867 | 0.78 | 742 | 0.54 | 643 | 0.46 |
| Reticulocytes | 827 | 0.64 | 676 | 0.41 | 575 | 0.37 |
| Lymphocytes | NA | NA | 651 | 0.35 | 642 | 0.43 |
| Hemoglobin | 1052 | 0.79 | 841 | 0.70 | 693 | 0.58 |
| MPV | 1373 | 0.91 | 1344 | 0.82 | 1094 | 0.73 |
| MCV | NA | NA | 746 | 0.56 | 676 | 0.55 |
| Monocytes | 875 | 0.80 | 638 | 0.64 | 708 | 0.67 |
| Platelets | NA | NA | 1247 | 0.59 | 1048 | 0.54 |
| PDW | 1037 | 0.83 | 1064 | 0.67 | 705 | 0.64 |
| RDW | NA | NA | 922 | 0.56 | 587 | 0.57 |
| RBCs | 1066 | 0.69 | 758 | 0.46 | 777 | 0.46 |
| BMD | 1400 | 0.78 | 1365 | 0.64 | 1079 | 0.57 |
| BMI | 1027 | 0.36 | 584 | 0.21 | 755 | 0.25 |
| Height | 2777 | 0.81 | 2740 | 0.63 | 2164 | 0.50 |
| DBP | 707 | 0.38 | 392 | 0.25 | 568 | 0.32 |
| SBP | 669 | 0.35 | 363 | 0.20 | 560 | 0.29 |
| College | NA | NA | 141 | 0.09 | 235 | 0.14 |
| HT | 391 | 0.24 | 180 | 0.13 | 299 | 0.22 |
| BMR | 1413 | 0.49 | 869 | 0.29 | 1080 | 0.36 |
| FEV1/FVC | 745 | 0.50 | 574 | 0.36 | 618 | 0.41 |
| FVC | 488 | 0.24 | 408 | 0.16 | 459 | 0.19 |
| Chronotype | NA | NA | 48 | 0.07 | 123 | 0.10 |

Table S1: Predicted vs. observed genome-wide significant results for 22 UK Biobank traits. We applied FMR and GENESIS to interim-release summary statistics (maximum N=145k) and predicted the number of significant loci and the heritability they would explain in the full release (maximum N=460k). We calculated observed values at N=460k using thresholding and pruning. For six traits, GENESIS predictions were NA because the estimated variance of the parameters was non-positive semidefinite (including two traits with negative variance estimates). GENESIS was run using the 3-component model and default settings. Projections were calculated at the exact full-release sample size for GENESIS (<460k) and at 460/145 times the interim-release sample size for FMR. MPV: mean platelet volume. MCV: mean cell volume: PDW: platelet distribution width. RDW: red cell distribution width. RBCs: red blood cell count. BMD: bone mineral density (heel). BMI: body mass index. DBP: diastolic blood pressure. SBP: systolic blood pressure. Edu: years of education. HT: hypertension. BMR: basal metabolic rate. FVC: forced vital capacity.

Table S2 (see Excel file): numerical results for Figures S2-S5.

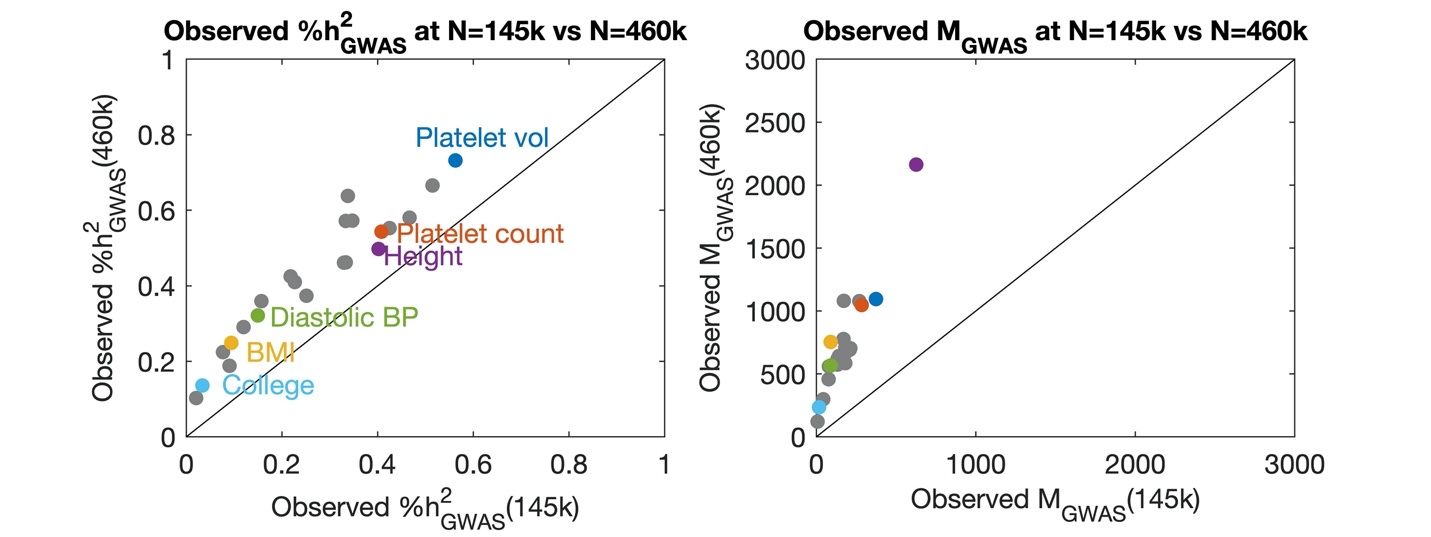

Figure S2: Observed number of genome-wide significant SNPs and proportion of heritability explained at N=145k vs 460k. For numerical results, see Table S2.

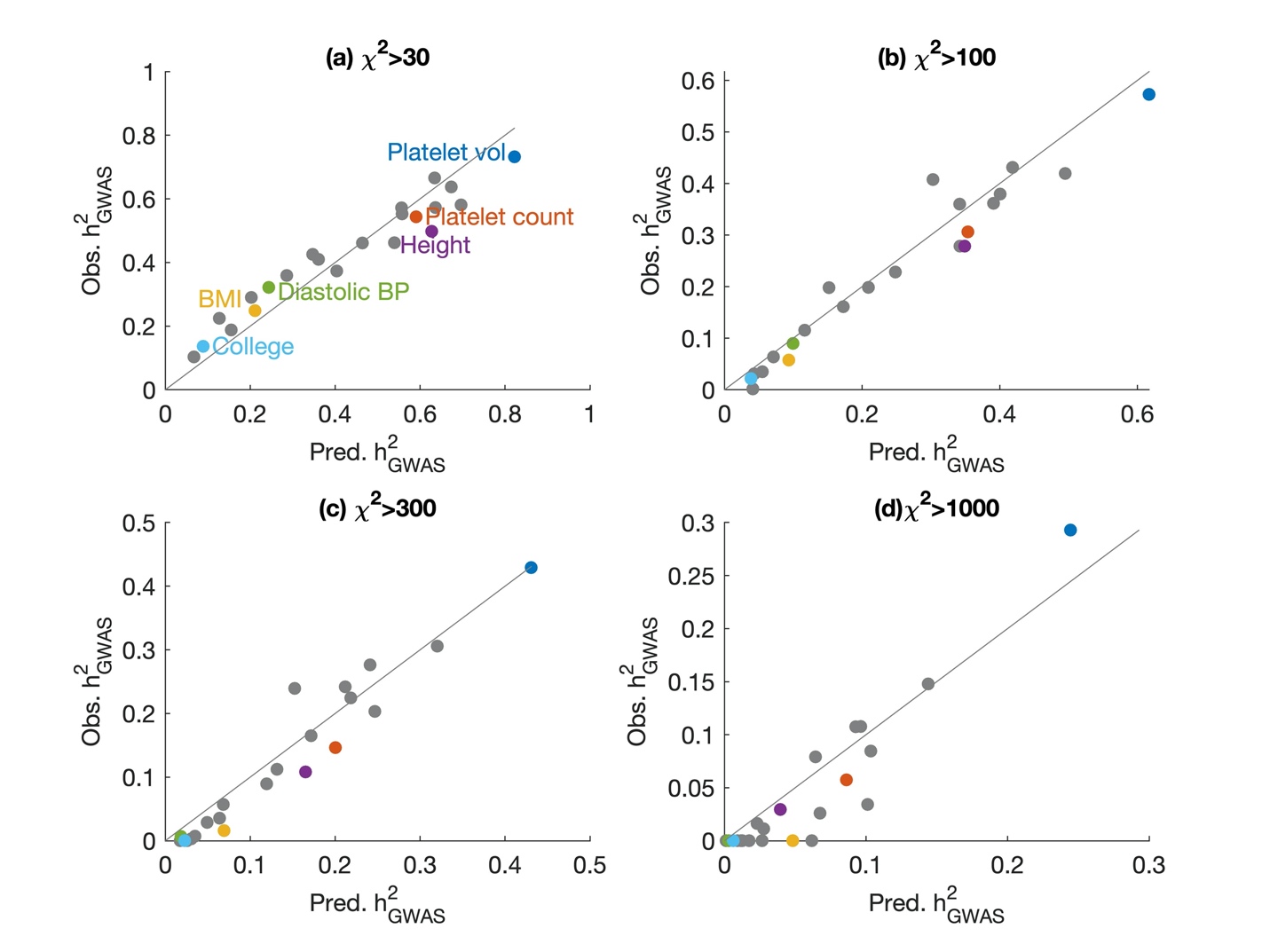

Figure S3: Predicted vs. observed heritability explained by genome-wide significant SNPs at different significance thresholds. %h2GWAS was predicted using interim-release UK Biobank summary statistics (maximum N=145k) and evaluated in the full release (maximum N=460k). For numerical results, see Table S2.

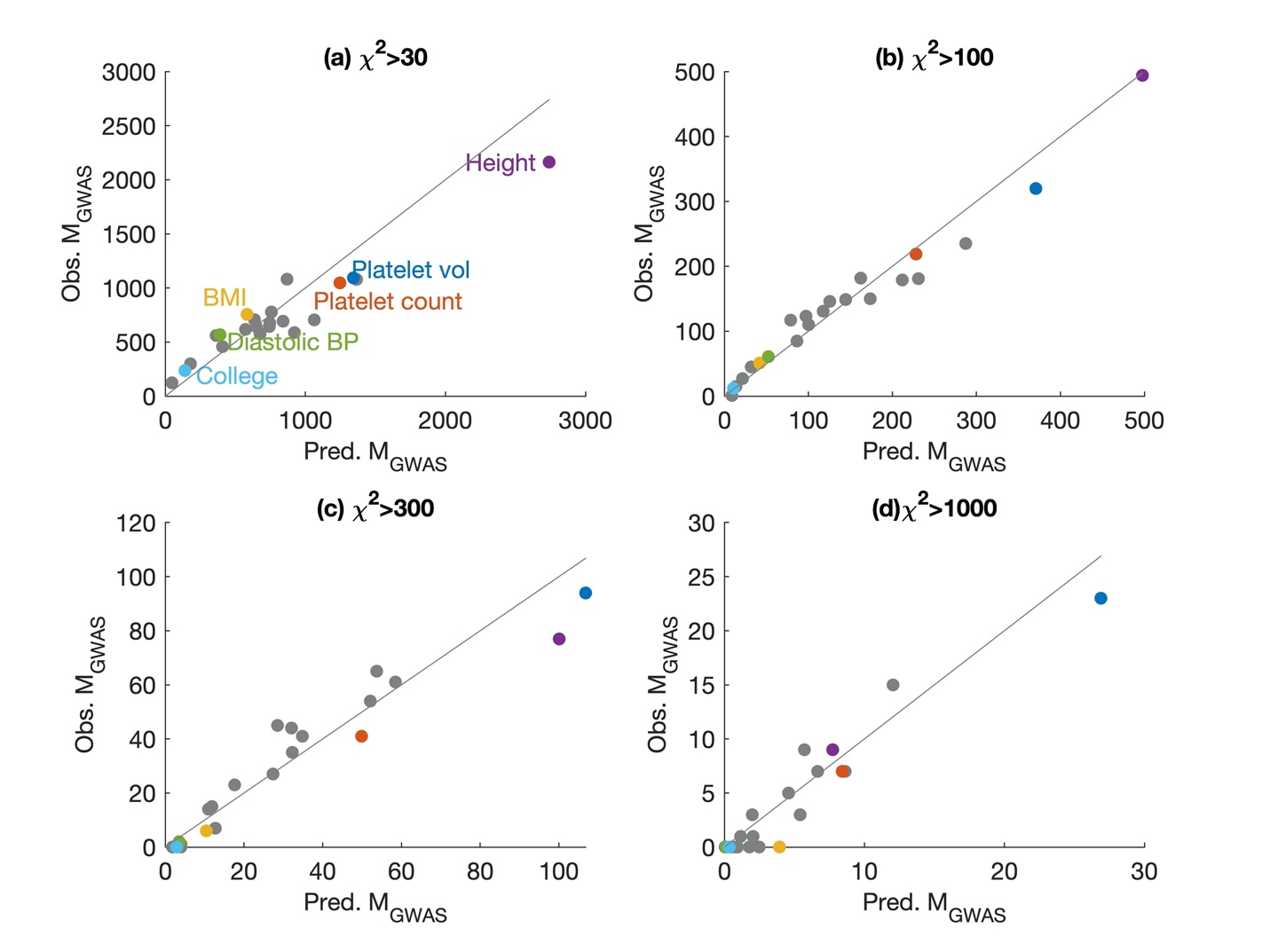

Figure S4: Predicted vs. observed number of genome-wide significant SNPs at different significance thresholds. MGWAS was predicted using interim-release UK Biobank summary statistics (maximum N=145k) and evaluated in the full release (maximum N=460k). For numerical results, see Table S2.

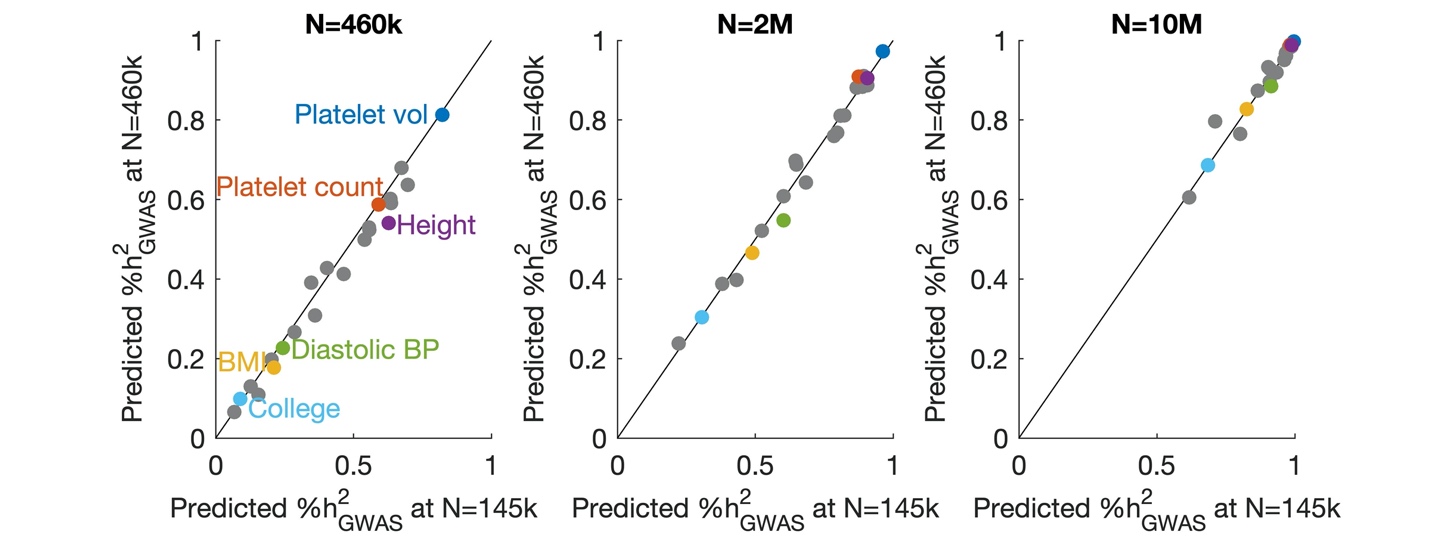

Figure S5: Predictions of %h2GWAS at large sample size, using N=145k vs. N=460k summary statistics. Estimates assume that the LD score regression intercept will be equal to what was observed at N=145k for both sets of estimates. For numerical results, see Table S2.

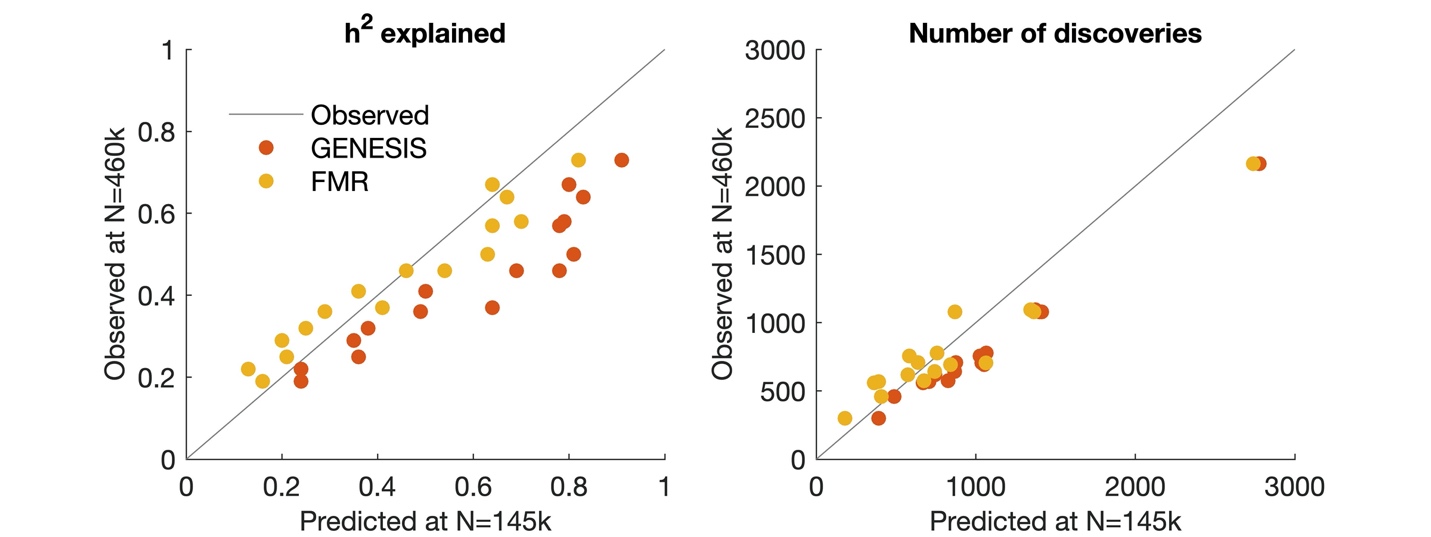

Figure S6: Performance of GENESIS vs. FMR predictions in UK Biobank. FMR and GENESIS were applied to interim-release UK Biobank summary statistics (maximum N=145k) in order to predict the results of the full release (maximum N=460k). Numerical results are presented in Table S1.

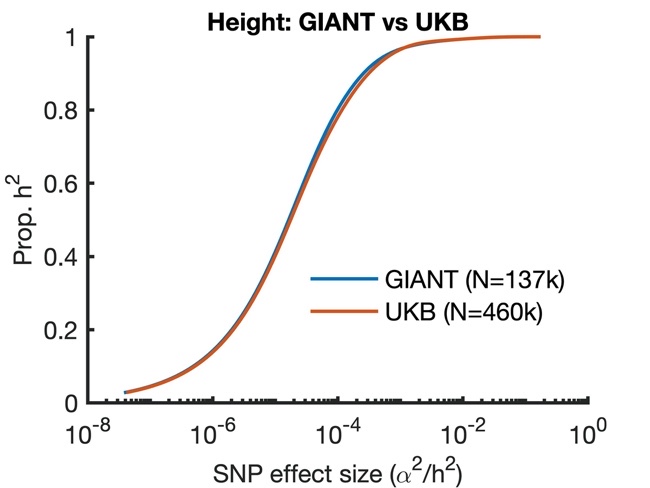

Figure S7: Estimated HDM of height using summary statistics from GIANT vs. UK Biobank. If results were biased by population stratification, we would expect the bottom-left portion of the curve (corresponding to small-effect SNPs) to be inflated for estimates based on GIANT.

Figure S8:
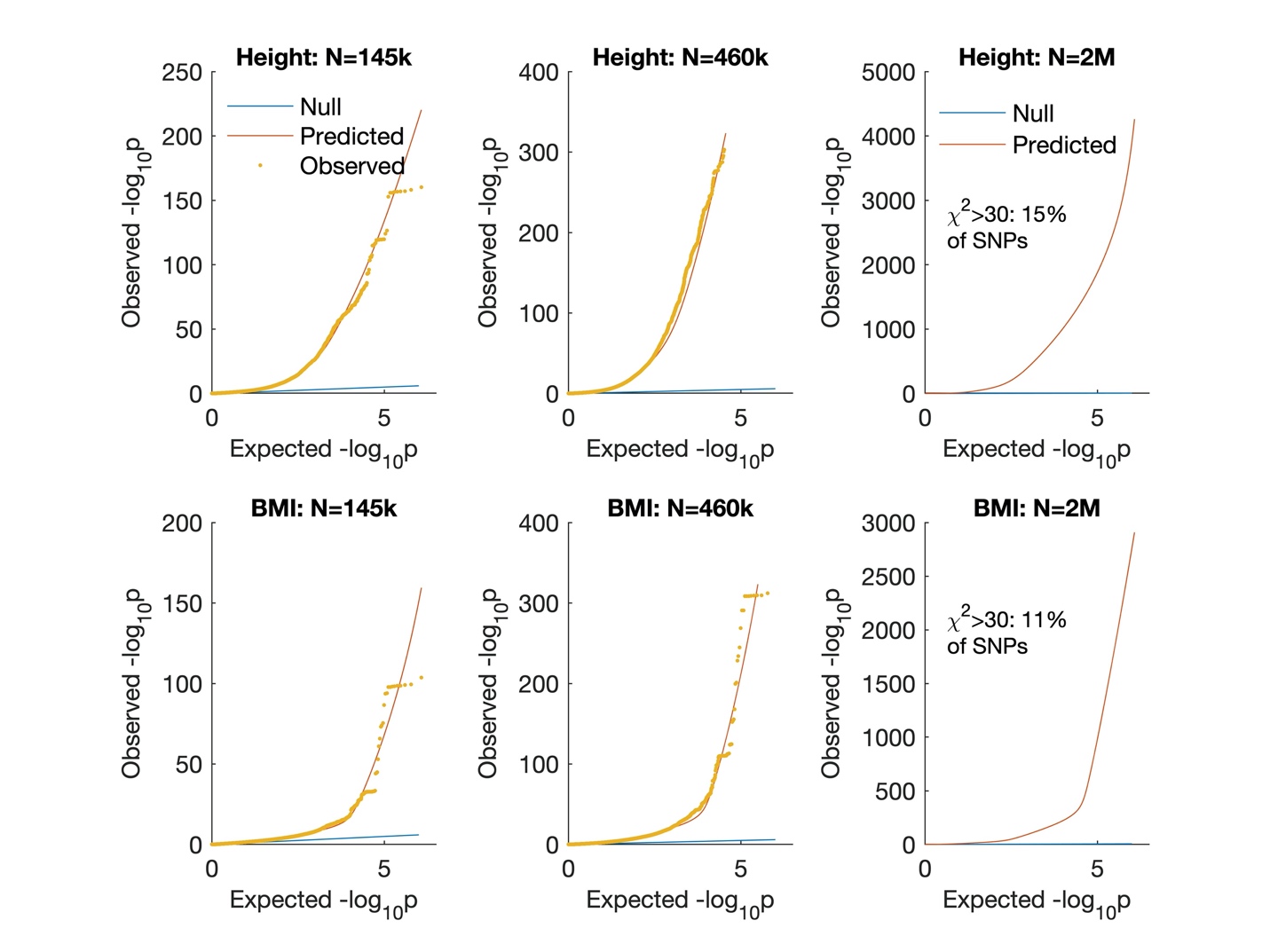
 Predicted vs. observed QQ plots for height and BMI. Predictions were generated by applying FMR-noLD to N=145k summary statistics.

|  | N_case_ | N_control_ | N_eff_ | N_tot_ |
| --- | --- | --- | --- | --- |
| Body mass index |  |  | 457824 | 457824 |
| Years of education |  |  | 454813 | 454813 |
| FVC (corrected for smoking) |  |  | 371949 | 371949 |
| Neuroticism | 234402 | 137664 | 346914 | 372066 |
| Blood pressure - diastolic |  |  | 422771 | 422771 |
| Height |  |  | 458303 | 458303 |
| Chronotype (morning person) |  |  | 410520 | 410520 |
| Age at menarche |  |  | 242278 | 242278 |
| Total protein level in blood |  |  | 397652 | 397652 |
| Waist-hip ratio (corrected for BMI) |  |  | 458417 | 458417 |
| FEV1/FVC (corrected for smoking) |  |  | 371949 | 371949 |
| Reaction time |  |  | 300486 | 300486 |
| Creatinine level |  |  | 434158 | 434158 |
| Schizophrenia | 40675 | 64643 | 99863 | 94437 |
| Platelet count |  |  | 444382 | 444382 |
| Sleep duration |  |  | 446118 | 446118 |
| IGF1 level in blood |  |  | 432292 | 432292 |
| General risk tolerance |  |  | 466571 | 466571 |
| Aspartate aminotransferase level |  |  | 430982 | 430982 |
| Red blood cell count |  |  | 442700 | 442700 |
| Insomnia | 108657 | 275291 | 311629 | 383948 |
| Atrial fibrillation | 60620 | 970216 | 228221 | 1030836 |
| Hypothyroidism | 22158 | 437166 | 84356 | 459324 |
| Allergy or eczema | 106051 | 352648 | 326129 | 458699 |
| Reported drinks per week |  |  | 527299 | 527299 |
| Phosphate levels |  |  | 397561 | 397561 |
| Breast cancer | 122977 | 105974 | 227688 | 228951 |
| Inflammatory bowel disease | 25042 | 34915 | 58331 | 59957 |
| Age at menopause |  |  | 143025 | 143025 |
| Bipolar disorder | 20352 | 31358 | 49367 | 46582 |
| Alzheimer’s disease | 71880 | 383378 | 242124 | 433886 |
| Coronary artery disease | 22233 | 64762 | 66204 | 86995 |

Table S3: 32 phenotypes used in main analyses. For the 11 binary phenotypes (10 diseases and neuroticism), we report case-control sample sizes, total sample sizes, and effective sample sizes (twice the harmonic mean). All summary statistics were publicly available (see URLs).

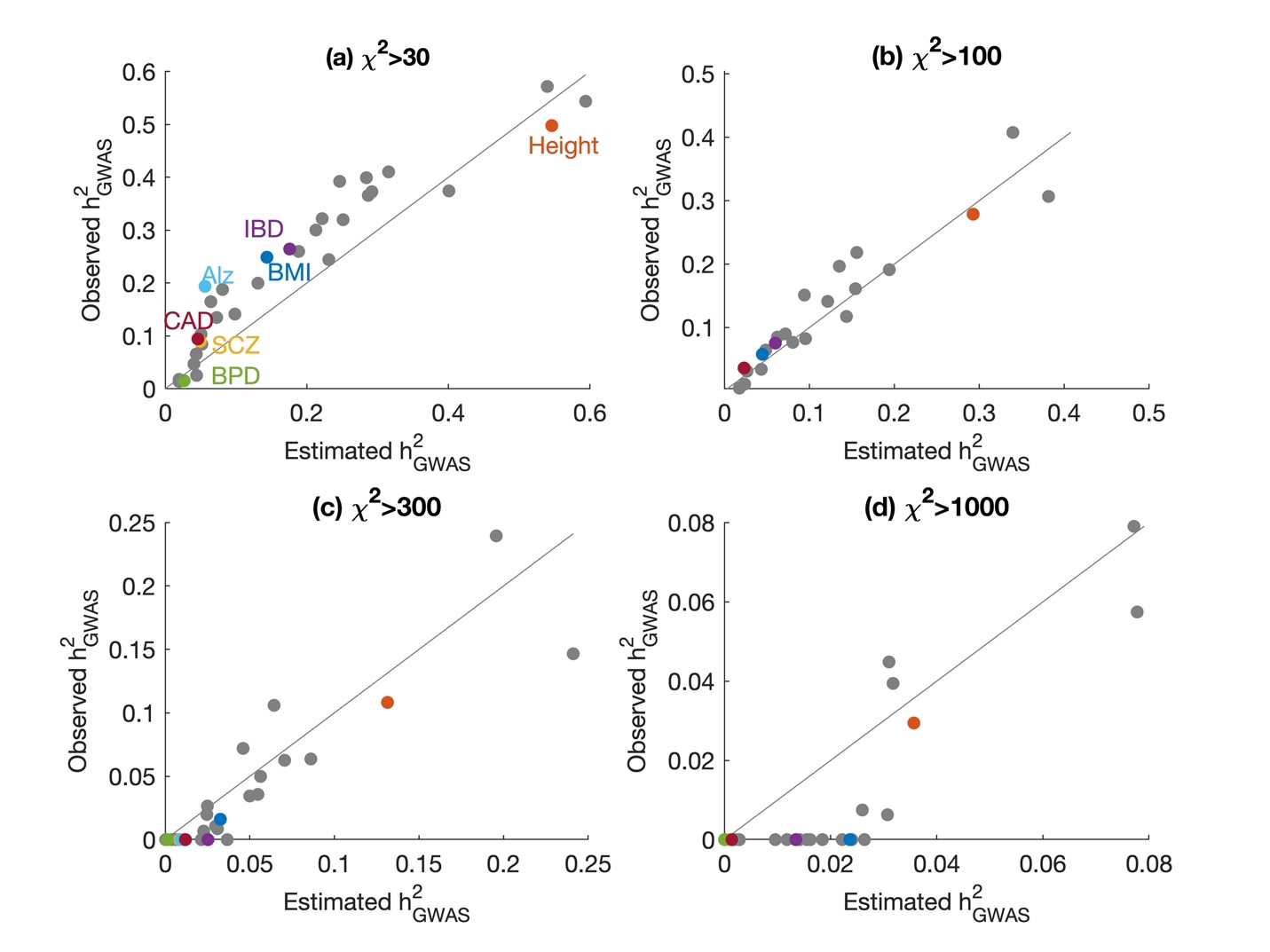

Figure S9: Estimated and observed %h2GWAS across 32 complex traits (Table S2) at four significance thresholds. $r^{2}$ was 0.90, 0.92, 0.83 and 0.73 in panels a-d respectively. For numerical results, see Table S4.

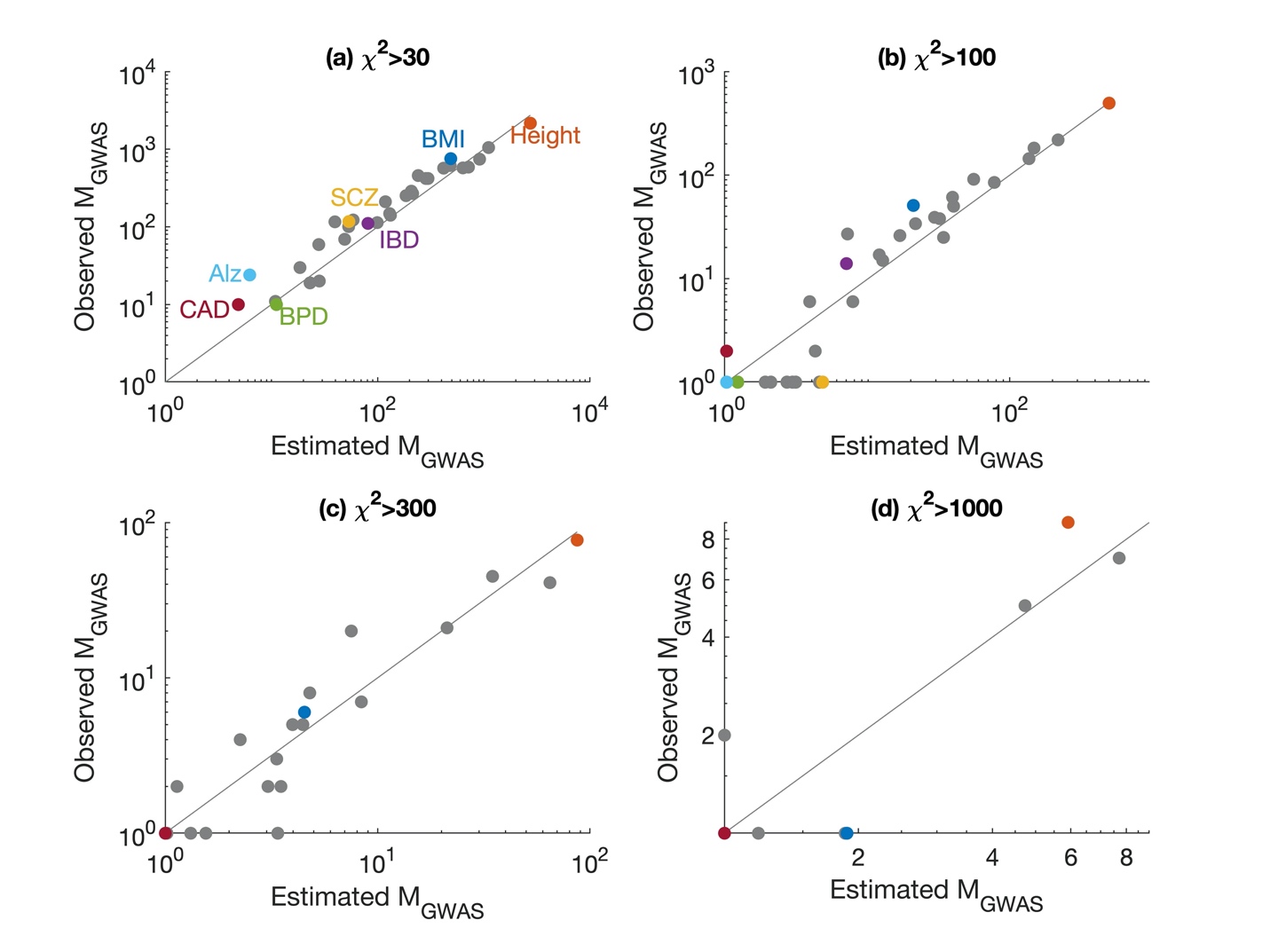

Figure S10: Estimated and observed number of GWS SNPs across 32 complex traits (Table S2) at four significance thresholds. Values below 1 are set to 1 for the purpose of plotting on a log scale; estimated and observed values are both smaller than 1 for 25/32 traits in panel (d). Linear-scale $r^{2}$ was 0.96, 0.99, 0.92and 0.87 in panels a-d respectively. For numerical results, see Table S4.

Table S4 (see Excel file): numerical results for Figures S9-10.

| Trait | N_eff_ (h2GWAS>0.5) | N_eff_  (h2GWAS >0.9) | Nh^2^ (h2GWAS>0.5) | Nh^2^ (h2GWAS>0.9) |
| --- | --- | --- | --- | --- |
| Body mass index | 2.9E+06 | 2.0E+07 | 1.1E+06 | 7.5E+06 |
| Years of education | 6.4E+06 | 5.1E+07 | 1.1E+06 | 8.4E+06 |
| FVC (corrected for smoking) | 3.0E+06 | 2.3E+07 | 9.6E+05 | 7.6E+06 |
| Neuroticism | 5.5E+06 | 4.9E+07 | 7.8E+05 | 6.9E+06 |
| Blood pressure - diastolic | 1.9E+06 | 1.3E+07 | 4.5E+05 | 3.2E+06 |
| Height | 4.1E+05 | 2.9E+06 | 3.5E+05 | 2.5E+06 |
| Chronotype (morning person) | 7.3E+06 | 5.8E+07 | 9.4E+05 | 7.5E+06 |
| Age at menarche | 1.9E+06 | 1.9E+07 | 6.8E+05 | 6.8E+06 |
| Total protein level in blood | 1.4E+06 | 8.9E+06 | 2.4E+05 | 1.5E+06 |
| Waist-hip ratio (corrected for BMI) | 2.3E+06 | 1.6E+07 | 4.0E+05 | 2.9E+06 |
| FEV1/FVC (corrected for smoking) | 1.0E+06 | 8.3E+06 | 2.9E+05 | 2.3E+06 |
| Reaction time | 7.5E+06 | 7.5E+07 | 6.1E+05 | 6.1E+06 |
| Creatinine level | 1.7E+06 | 1.7E+07 | 5.7E+05 | 5.7E+06 |
| Schizophrenia | 1.6E+06 | 1.3E+07 | 8.4E+05 | 6.7E+06 |
| Platelet count | 2.8E+05 | 2.5E+06 | 1.1E+05 | 9.8E+05 |
| Sleep duration | 8.9E+06 | 1.0E+08 | 6.8E+05 | 7.6E+06 |
| IGF1 level in blood | 7.7E+05 | 7.7E+06 | 2.8E+05 | 2.8E+06 |
| General risk tolerance | 1.5E+07 | 2.1E+08 | 1.6E+06 | 2.2E+07 |
| Aspartate aminotransferase level | 1.5E+06 | 1.1E+07 | 1.8E+05 | 1.3E+06 |
| Red blood cell count | 3.9E+05 | 4.4E+06 | 1.1E+05 | 1.2E+06 |
| Insomnia | 1.4E+07 | 2.2E+08 | 1.0E+06 | 1.6E+07 |
| Atrial fibrillation | 3.6E+06 | 4.1E+07 | 4.7E+05 | 5.2E+06 |
| Hypothyroidism | 5.3E+05 | 6.0E+06 | 1.7E+05 | 1.9E+06 |
| Allergy or eczema | 2.3E+06 | 2.3E+07 | 2.7E+05 | 2.7E+06 |
| Reported drinks per week | 1.3E+07 | 1.1E+08 | 8.1E+05 | 6.4E+06 |
| Phosphate levels | 2.0E+06 | 1.8E+07 | 2.7E+05 | 2.4E+06 |
| Breast cancer | 9.1E+05 | 8.1E+06 | 1.3E+05 | 1.2E+06 |
| Inflammatory bowel disease | 3.3E+05 | 3.3E+06 | 1.4E+05 | 1.4E+06 |
| Age at menopause | 2.3E+06 | 2.5E+07 | 4.6E+05 | 5.2E+06 |
| Bipolar disorder | 2.2E+06 | 2.8E+07 | 9.4E+05 | 1.2E+07 |
| Alzheimer’s disease | 1.4E+07 | 1.4E+08 | 3.2E+05 | 3.2E+06 |
| Coronary artery disease | 1.7E+06 | 1.7E+07 | 1.6E+05 | 1.6E+06 |

Table S5: Sample-size targets across 32 diseases and complex traits, estimated using FMR. We report the estimated sample size (or effective sample size) required for genome-wide significant SNPs to explain 50% or 90% of SNP-heritability, and the effective signal strength (Nh^2^_eff_) required for genome-wide significant SNPs to explain 50% or 90% of SNP-heritability. For case-control traits, if there are a large number of controls, the required number of cases is ¼ the required N_eff_.

Table S6 (see Excel file): Numerical results for Figures 3 and S11. We also report $Nh_{eff}^{2}$ targets and the number of loci that will be associated when 90% of heritability is explained.

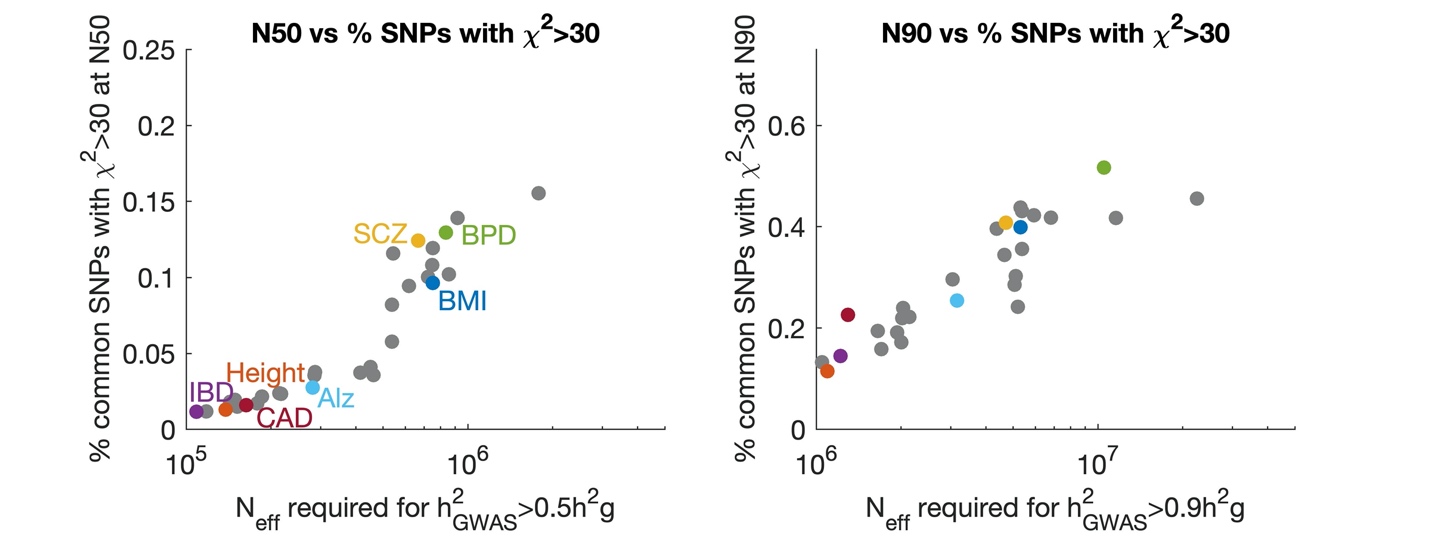

Figure S11: Proportion of genome-wide significant SNPs when %h2GWAS exceeds 0.5 or 0.9. This proportion is highly correlated with the observed signal that is required to reach the %h2GWAS threshold, because both are higher for highly polygenic traits. Numerical results are presented in Table S6.

| Trait | %h2GWAS | MGWAS | r^2^_PGS_/h^2^_g_ (Loh et al 2018) | MGWAS (Loh et al. 2018) |
| --- | --- | --- | --- | --- |
| Height | 0.50 | 2164 | 0.75 | 2098 |
| FEV1/FVC (corrected for smoking) | 0.41 | 618 | 0.49 | 566 |
| Platelet count | 0.54 | 1048 | 0.7 | 1007 |
| Red blood cell count | 0.57 | 587 | 0.58 | 714 |

Table S7: Previously-reported PGS accuracy for well-powered UK Biobank traits. Loh et al (2018) reported PGS r2 using BOLT-LMM for these four traits using the exact same data, and the reported number of significant loci approximately agreed with our thresholding-and-pruning approach. We derived an upper bound on PGS r^2^ that is expected to be tight for an optimal PGS method at adequate sample size; this bound is close to 0.9 times the SNP-heritability when %h2GWS is approximately 0.5. Loh et al. reported smaller PGS r^2^ values.

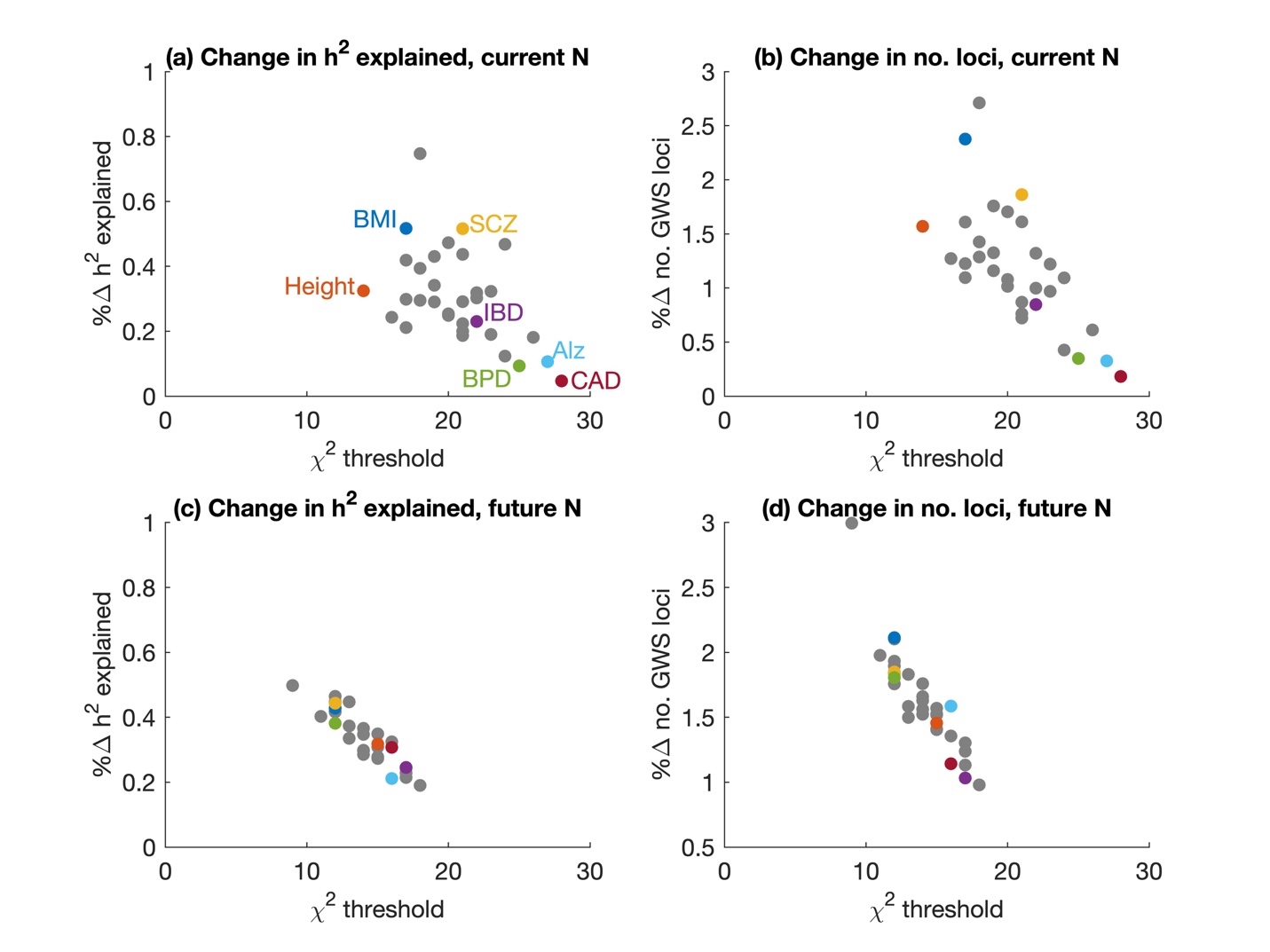

Figure S12: Effect of calibrating the GWAS significance threshold to the NTPR. We report the $\chi^{2}$ threshold that corresponds to an estimated NTPR of 99% and the change in heritability explained and number of significant loci at this threshold instead of genome-wide significance ($\chi^{2}>30)$. (a) $\chi^{2}$ threshold that corresponds to an estimated NTPR of 99% at current sample size, and percent change in heritability explained by SNPs exceeding this threshold. Heritability explained was calculated from FMR estimates instead of using pruning and thresholding, as the latter is expected to suffer from substantial bias due to winner’s curse at $\chi^{2}$ thresholds well below 30. (b) Change in the number of significant loci at current sample size, calculated from FMR estimates. (c) $\chi^{2}$ threshold that corresponds to an estimated NTPR of 99% at the sample size where significant SNPs are predicted to explain 50% of heritability, and percent change in heritability explained by SNPs exceeding this threshold (e.g. a percent change of 50% corresponds to a total of 75%). (d) Change in the number of significant loci at the sample size where significant SNPs are predicted to explain 50% of heritability. Numerical results are presented in Table S9.

Table S8 (see Excel file): numerical results for Figure 4. We also report how NTPR estimates change when different values of $M_{\mathrm{harmonic}}$ are assumed (see Methods).

Table S9 (see Excel file): numerical results for Figure S12. “Future N” refers to the sample size at which GWS SNPs explain 50% of heritability (which is already the case for a few traits).

Table S10 (see Excel file): quantiles of heritability for each trait, with jackknife standard errors. We also report meta-analyzed values, and the values that are expected under the log-normal model fit.

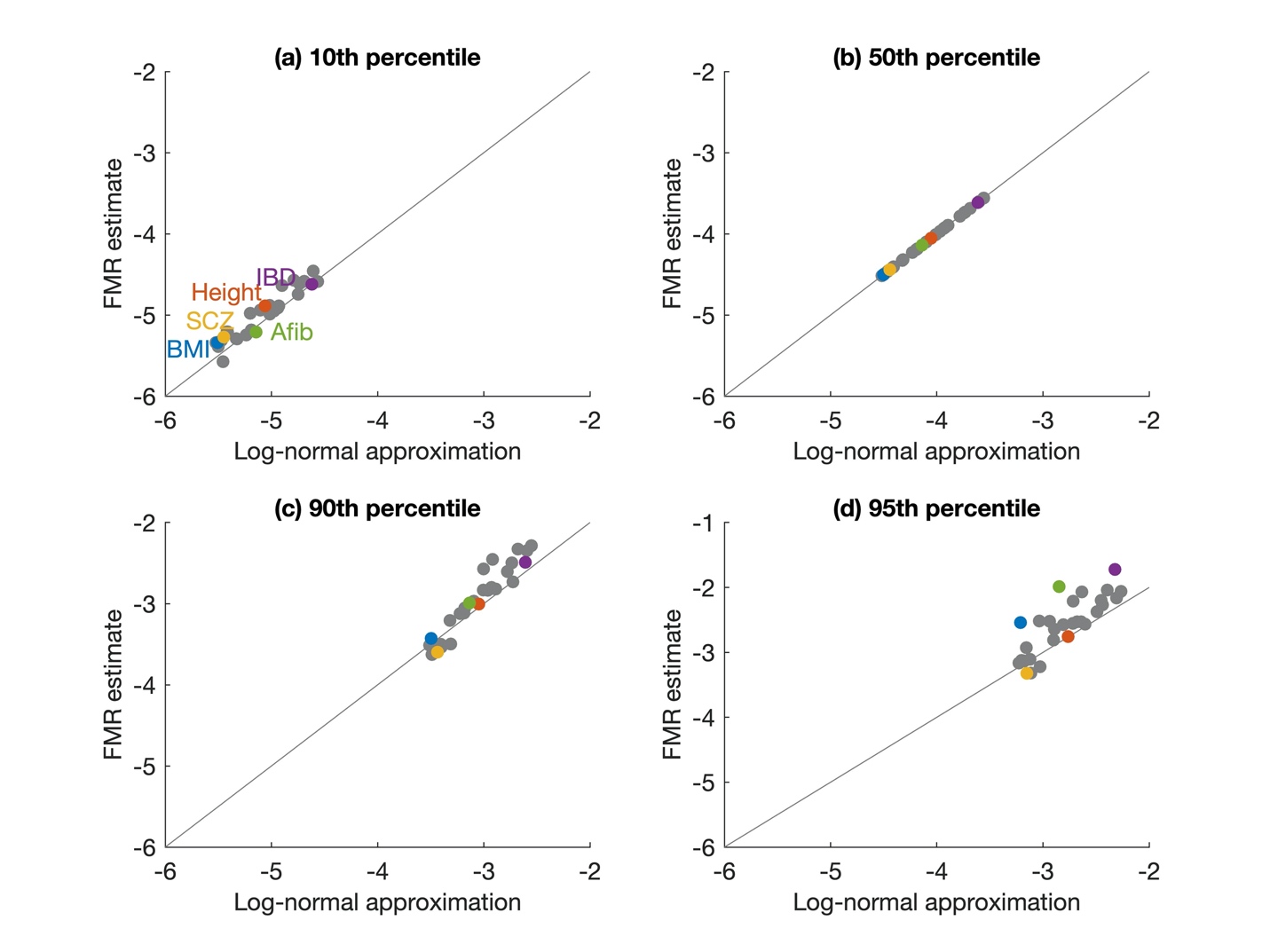

Figure S13: Fit of the log-normal model. Quantiles are plotted for the FMR estimate vs. the log-normal approximation for well-powered traits (same criterion as Figure 5), on a log-10 scale.

Numerical values are presented in Table S10.
